# Simultaneous inhibition of human CD4 and 4-1BB biogenesis suppresses cytotoxic T lymphocyte proliferation

**DOI:** 10.1101/2020.12.30.424816

**Authors:** Elisa Claeys, Eva Pauwels, Stephanie Humblet-Baron, Dominique Schols, Mark Waer, Ben Sprangers, Kurt Vermeire

## Abstract

The small molecule cyclotriazadisulfonamide (CADA) down-modulates the human CD4 receptor, an important factor in T cell activation. Here, we addressed the immunosuppressive potential of CADA using *in vitro* activation models. CADA inhibited lymphocyte proliferation in a mixed lymphocyte reaction, and when human PBMCs were stimulated with CD3/CD28 beads or phytohemagglutinin. The immunosuppressive effect of CADA involved both CD4^+^ and CD8^+^ T cells but was, surprisingly, most prominent in the CD8^+^ T cell subpopulation where it inhibited cell-mediated lympholysis. We discovered a direct down-modulatory effect of CADA on 4-1BB (CD137) expression, a survival factor for activated CD8^+^ T cells. More specifically, CADA blocked 4-1BB protein biosynthesis by inhibition of its co-translational translocation across the ER membrane in a signal peptide-dependent way. This study demonstrates that CADA, as potent down-modulator of human CD4 and 4-1BB, has promising *in vitro* immunomodulatory characteristics for future *in vivo* exploration as immunosuppressive drug.

## INTRODUCTION

The cluster of differentiation 4 (CD4) receptor is a type I integral membrane protein consisting of four extracellular immunoglobulin-like domains, a spanning transmembrane region and a short cytoplasmic tail (Maddon et al., 1985). The lymphocyte C-terminal Src kinase (Lck) non-covalently interacts with the cytoplasmic tail of CD4 (Shaw et al., 1989). Next to its function in CD4 signaling, Lck inhibits endocytosis of the CD4 receptor by preventing the entry of CD4 into clathrin-coated pits (Pelchen-Matthews et al., 1992). Several immune cell types express the CD4 receptor with T helper cells expressing the highest levels, followed by monocytes that express already 10- to 20-fold less CD4 compared to T cells (Collman et al., 1990). Studies in CD4 null mice underline the role of the CD4 receptor in positive thymic selection and development of helper T cells (Rahemtulla et al., 1991).

The CD4 receptor is also crucial for proper immune function, especially during T cell activation in which it can fulfil several roles (Claeys et al., 2019). The CD4 receptor can exert an intercellular adhesion function by stabilizing the interaction between the T cell receptor on CD4^+^ T cells and the major histocompatibility complex class II on antigen-presenting cells (Janeway, 1989). More important are the signaling function of the CD4 receptor in T cell activation through Lck and the enhancement of T cell sensitivity to antigens mediated by CD4 (Konig et al., 2004; Krogsgaard et al., 2005). Besides its role in T cell activation, the CD4 receptor is suggested to be involved in peripheral T cell differentiation towards the T helper 2 subset and in the chemotactic response of CD4^+^ T cells towards interleukin (IL)-16 (Cruikshank et al., 1991; Fowell et al., 1997). Additionally, different functions are attributed to the CD4 receptor in other types of immune cells including natural killer and dendritic cells (Bernstein et al., 2006; Bialecki et al., 2011). The important role of the CD4 receptor in the immune system has been further demonstrated by the *in vitro* and *in vivo* immunosuppressive potential of non-depleting anti-CD4 monoclonal antibodies (Mayer et al., 2013; Schulze-Koops et al., 2000; Winsor-Hines et al., 2004).

In the field of virology, attachment of viral gp120 of human immunodeficiency virus (HIV) to the cellular CD4 receptor initiates HIV infection of target cells (Dalgleish et al., 1984; Klatzmann et al., 1984). From an antiviral screen, the small molecule cyclotriazadisulfonamide (CADA) was identified as a potent inhibitor of HIV infection (Vermeire et al., 2002). The antiviral effect of this synthetic macrocycle is due to down-modulation of the CD4 protein, the primary entry receptor for HIV (Vermeire et al., 2003). This down-modulating activity of CADA is reversible *in vitro*: when treatment is ceased, cellular CD4 expression is rapidly restored to normal levels (Vermeire et al., 2007). Additionally, CADA does not compromise cellular viability as was demonstrated by long-term (about 1 year) exposure of a T cell line to CADA, with full recovery of CD4 expression when treatment was terminated (Vermeire et al., 2014). The sensitivity of the CD4 receptor to CADA is species-specific, as expression of murine CD4 (mCD4) was not affected by CADA, while primary T cells of macaques responded in a similar way as human T cells. Mechanistically, CADA was shown to inhibit endoplasmic reticulum (ER) co-translational translocation of the human CD4 (hCD4) pre-protein in a signal peptide (SP)-dependent way (Vermeire et al., 2014). CADA selectively binds to the SP of hCD4, thereby locking it in an intermediate conformation inside the Sec61 translocon channel during co-translational translocation through the ER membrane, finally resulting in proteasomal degradation in the cytosol of the mistranslocated hCD4 precursor molecules. The CADA-sensitive region of hCD4 consists primarily of the hydrophobic core of the hCD4 SP, although the presence of charged residues in the N-terminal portion of the mature protein enhances sensitivity (Van Puyenbroeck et al., 2020). Almost no binding of CADA to the mCD4 SP was detected, explaining the observed resistance of mCD4 to CADA (Vermeire et al., 2014).

Thus, CADA down-modulates the CD4 receptor, a key component in T cell activation. Therefore, we explored in this study whether CADA has a potential immunomodulatory capacity. Here, CADA was evaluated in several *in vitro* models of T cell activation and was found to exert a clear immunosuppressive effect. Furthermore, in addition to the earlier reported CD4 receptor, we identified 4-1BB – a crucial co-stimulatory factor in T cell activation of mainly cytotoxic lymphocytes – as a new target of CADA.

## RESULTS

### CADA down-modulates the human CD4 receptor and has an immunosuppressive effect in the mixed lymphocyte reaction

In line with our previous report (Vermeire et al., 2014), the small molecule CADA (***Figure 1A***) dose-dependently down-modulated the hCD4 receptor on Jurkat T cells as well as on human peripheral blood mononuclear cells (PBMCs) (***Figure 1B***). At a concentration of 10 μM CADA, the cell surface hCD4 expression level was greatly reduced in both cell types: 86% reduction in hCD4 expression for Jurkat cells and 74% for PBMCs, as compared to untreated control cells (IC_50_ values of 0.41 μM and 0.94 μM, respectively). Based on this hCD4 receptor down-modulating potency of CADA, we addressed whether CADA has a potential immunomodulatory capacity in human cells. In a first approach, the effect of CADA was evaluated in T cells activated *in vitro* by means of superantigens. Limited or no inhibitory effect of CADA on the expression of the early activation marker CD69 was observed when Jurkat T cells were activated by the superantigen staphylococcal enterotoxin E (SEE), nor when naive CD4^+^ T cells were activated by SEE or staphylococcal enterotoxin B (SEB) (***Figure 1–figure supplement 1***). However, CADA significantly inhibited lymphocyte proliferation in the mixed lymphocyte reaction (MLR) in which PBMCs are co-cultured with mitomycin-inactivated stimulator B cells (***Figure 1C***). Although lymphocyte proliferation was not blocked completely, there was a strong dose-dependent inhibitory effect of CADA. The antiproliferative immunosuppressive agent mycophenolate mofetil (MMF), included as control, evoked a stronger maximal inhibitory effect, with complete inhibition of lymphocyte proliferation at a dose of 2 μM of MMF and higher (***Figure 1C***). Viability of Jurkat cells cultured in the presence of CADA was not affected for concentrations up to 50 μM as determined by trypan blue staining (***Figure 1–figure supplement 2***), and only a small reduction in metabolic activity (as quantified by MTS-PES) was observed for higher doses of CADA that reached significance at a concentration of 50 μM (***Figure 1D***). In contrast, a reduction in cell viability (***Figure 1–figure supplement 2***) and a significant dose-dependent inhibition of metabolic activity was observed for cells treated with MMF (***Figure 1D***).

**Figure 1.**
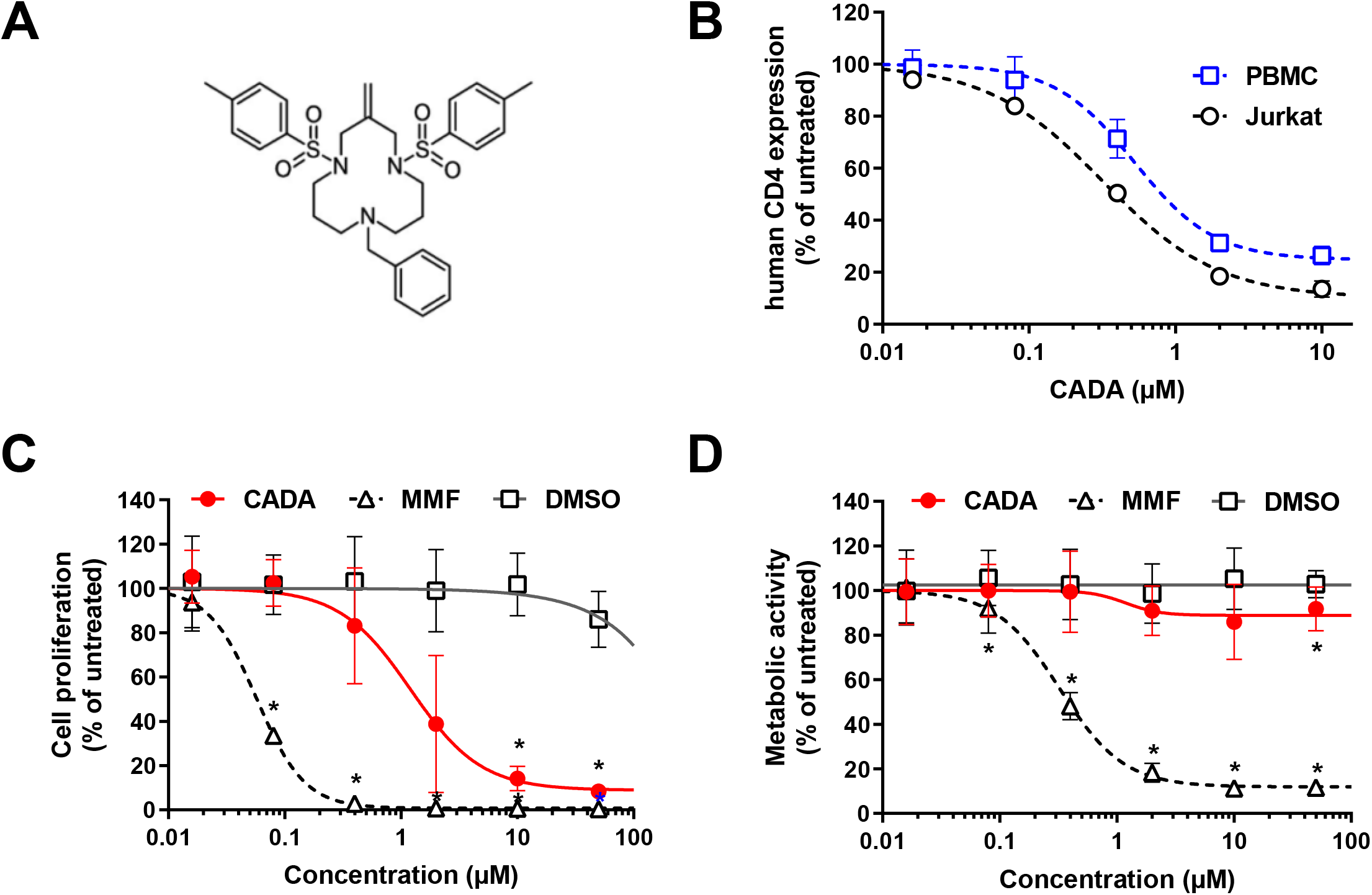
CADA down-modulates the human CD4 receptor and has an immunosuppressive effect in the mixed lymphocyte reaction. **(A)** Chemical structure of cyclotriazadisulfonamide or CADA (9-benzyl-3-methylene-1,5-di-*p*-toluenesulfonyl-1,5,9-triazacyclododecane). **(B)** Four parameter dose-response curves for CADA of cell surface human CD4. Cells were incubated with increasing concentrations of CADA and CD4 expression was measured by flow cytometry using a PE-labeled anti-human CD4 antibody (clone SK3) after 2 days for Jurkat cells (n=3) or 5 days for PBMCs (n=3). CD4 expression is given as percentage of untreated control (mean ± SD). **(C)** PBMCs were co-cultured with mitomycin C inactivated RPMI1788 cells in the presence of CADA, MMF or matching DMSO concentrations. At day 5, [^3^H]-thymidine was added and proliferation response was measured 18h later by detecting counts per minute. Lymphocyte proliferation is given as percentage of untreated control (mean ± SD; n=4). Multiple t-tests were performed to compare each concentration of CADA or MMF to the corresponding DMSO control with *p<0.05 and with Holm-Sidak method as correction for multiple comparison. **(D)** Jurkat cells were exposed to different concentrations of CADA, MMF or DMSO during 2 days, after which MTS-PES was added to measure cellular metabolic activity, and read-out was done 2h later on a spectrophotometer. Metabolic activity of cells is given as percentage of untreated control (mean ± SD; n=10). Multiple t-tests were performed to compare each concentration of CADA or MMF to the corresponding DMSO control with *p<0.05 and with Holm-Sidak method as correction for multiple comparison. **Figure supplement 1.** CADA has no effect on superantigen-activated lymphocytes. **Figure supplement 2.** CADA does not exert cytotoxicity.

### Reduced CD4 surface expression affects lymphocyte proliferation in the MLR

As CADA down-modulates the hCD4 receptor, we next investigated if reduced cell surface CD4 expression correlates with inhibition of lymphocyte proliferation. Therefore, we compared CADA with another agent that directly targets the hCD4 receptor, namely the non-depleting anti-CD4 monoclonal antibody Clenoliximab (Hepburn et al., 2003). PBMCs were co-cultured with mitomycin-inactivated RPMI1788 cells in the presence of the compound, and at day five, the sample was evaluated for CD4 expression by flow cytometry. In parallel, an identical sample was exposed to [^3^H]-thymidine to measure the proliferation response 18h later. CADA-treatment resulted in a consistent dose-dependent reduction in CD4 expression, that reached a plateau at 2 μM of CADA (***Figure 2***, left panel). Treatment with Clenoliximab also had a CD4 down-modulating effect but this was less effective and more variable as compared to CADA (***Figure 2***, right panel). In addition, there was an inhibitory effect of Clenoliximab seen on lymphocyte proliferation, although rather limited (about 30% reduction) and less evident as the reduction in CD4 expression (***Figure 2***). For CADA, a clear dose-dependent inhibition of lymphocyte proliferation was observed (***Figure 2***, left panel). However, whereas CD4 reduction plateaued at 2 μM of CADA, a steady decrease in lymphocyte proliferation was measured with increasing doses of CADA. This suggests that for CADA (an) additional immunomodulatory effect(s) are at play beyond suppression of CD4 receptor expression.

**Figure 2.**
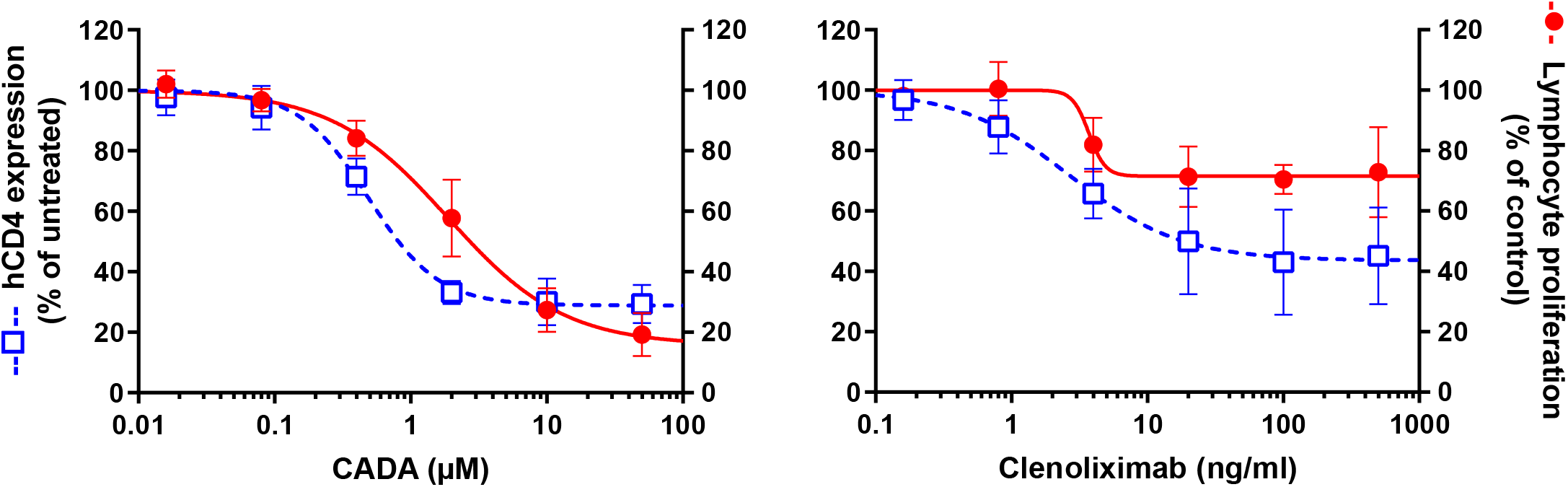
Reduced CD4 surface expression affects lymphocyte proliferation in the MLR. PBMCs were co-cultured with mitomycin C inactivated RPMI1788 cells in the presence of CADA (left panel) or the anti-CD4 antibody Clenoliximab (right panel). At day 5, one sample was used to determine cell surface human CD4 expression using flow cytometry. In parallel, [^3^H]-thymidine was added to an identical sample and proliferation response was measured by detecting counts per minute 18h later. To avoid steric hindrance for the detection of CD4, the monoclonal anti-human CD4 antibody clone OKT4 was used as this antibody binds to the D3 domain of CD4, while Clenoliximab binds to the D1 domain. Human CD4 expression (open blue symbols with dotted line), given as percentage of untreated control, is plotted on the left Y-axis (mean ± SD; n=4), and lymphocyte proliferation (solid red symbols with solid line), given as percentage of DMSO control for CADA and as percentage of ProClin 300 control for Clenoliximab is plotted on the right Y-axis (mean ± SD; n=4).

### CADA suppresses lymphocyte proliferation and inhibits upregulation of CD4 and CD8 after activation by CD3/CD28 beads or PHA

To further explore the inhibitory effect of CADA on lymphocyte proliferation, we evaluated CADA in two additional *in vitro* models of T cell activation. The first one, referred to as CD3/CD28 beads stimulation assay, is based on the use of inert, superparamagnetic beads to which anti-CD3 and anti-CD28 antibodies are covalently coupled. The second model is by addition of phytohemagglutinin (PHA), a lectin that binds to sugars on glycosylated surface proteins, including the TCR and CD3, thereby crosslinking them. Briefly, PBMCs were pre-incubated with a fixed dose of CADA (10 μM) or DMSO control for 3 days before activation by CD3/CD28 beads or PHA. In both models, the proliferation response of lymphocytes in the control samples steadily increased over time in all donors, with a peak at day 3 post activation (***Figure 3***; open symbols). Treatment with CADA suppressed the responsiveness of lymphocytes to both CD3/CD28 beads and PHA (***Figure 3***; solid red symbols). Intra-donor analysis revealed that CADA significantly reduced cell proliferation compared to DMSO control in both models at day 1 and 2 post activation, as further exemplified by the insert panels of ***Figure 3*** (p = 0.002 and p = 0.003 for CD3/CD28 and PHA, respectively; paired t-test).

**Figure 3.**
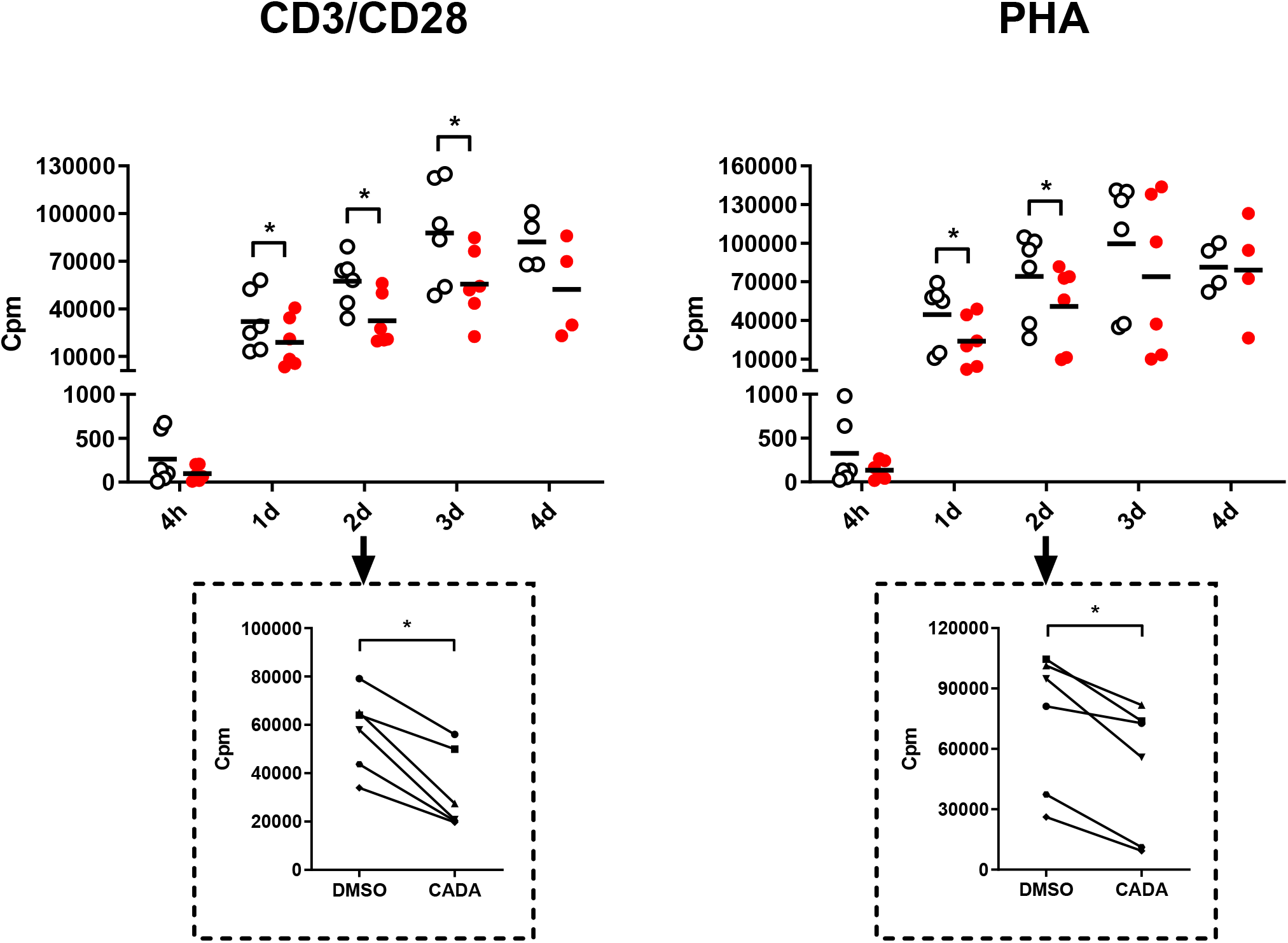
CADA suppresses lymphocyte proliferation after activation by CD3/CD28 beads or PHA. PBMCs were pre-incubated with CADA (10 μM) or DMSO during 3 days, after which they were activated by CD3/CD28 beads (left panels) or PHA (right panels). At 4h, 1d, 2d, 3d or 4d post activation, [3H]-thymidine was added and proliferation response was measured by detecting counts per minute (cpm) 22h later. Individual cpm values are shown for stimulated PBMCs with DMSO-treated cells as open symbols and CADA-treated cells as solid red dots. Horizontal lines indicate the mean values of 4 to 6 donors. Insert panels below the graph show intra-donor treatment effect on the proliferation response at day 2 post activation (each donor is indicated separately). A paired t-test was performed to compare CADA to DMSO with *p<0.05.

In addition to the proliferation response, we analyzed the expression level of cell surface CD4 and CD8, receptors known to be involved in T cell activation. As expected, basal CD4 expression on CD4^+^ T cells measured at time point 0, which is after 3 days of CADA pre-incubation, was decreased by half in the CADA-treated samples (***Figure 4A***, ***Figure 4–figure supplement 1***). In control CD4^+^ T cells (treated with DMSO) cell surface CD4 expression was strongly upregulated starting from day 1 post activation by CD3/CD28 beads and by PHA (***Figure 4A***). In sharp contrast, in both activation models CADA completely blocked this induced CD4 upregulation in all donors and at every tested time point (***Figure 4A***, ***Figure 4–figure supplement 1***), a result of the complete inhibition of hCD4 protein biogenesis by CADA (Vermeire et al., 2014). In the CD8^+^ T cell population, basal CD8 expression was also partially affected by pre-treatment with CADA (***Figure 4B***, ***Figure 4–figure supplement 1***). Intra-donor flow cytometric analysis of the samples revealed that the mean fluorescence intensity (MFI) for CD8 receptor expression in the CADA-treated cells was reduced by 38 ± 4% (mean ± SD; ***Figure 4–figure supplement 1***, d0). After activation by CD3/CD28 beads and PHA, CD8 expression was upregulated in the control samples, starting at day 1 and with a continuous increase over the next days. Exposure of the cells to CADA clearly suppressed this activation-triggered CD8 upregulation (***Figure 4B***). However, from day 3 onwards, CD8 levels started to rise in the CADA-treated samples, which was most prominent in the PHA-stimulated cells (***Figure 4B***, right panel). Consequently, the suppression of CADA on CD8 receptor upregulation in these cells plateaued around 50% (***Figure 4–figure supplement 1***).

**Figure 4.**
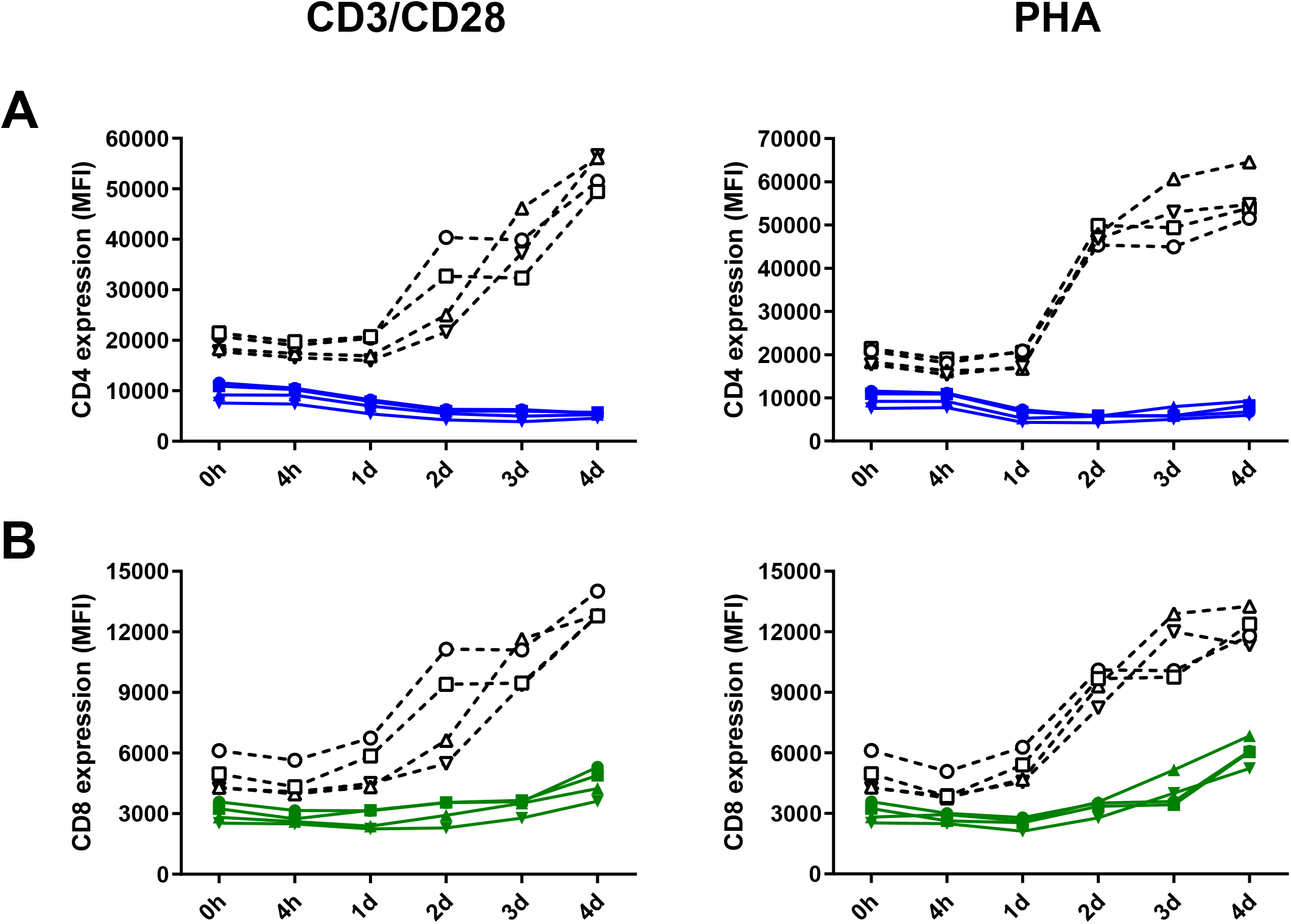
CADA inhibits upregulation of CD4 and CD8 after activation by CD3/CD28 beads or PHA. **(A and B)** Cell surface CD4 (A) and CD8 (B) receptor expression was measured by flow cytometry just before activation (0h) and 4h, 1d, 2d, 3d or 4d post activation with CD3/CD28 (left) or PHA (right). Mean fluorescence intensity (MFI) of human CD4 or CD8 receptor expression is shown for 4 donors of PBMCs (indicated separately) with DMSO-treated samples as a dotted line and CADA-treated samples as a full line. **Figure supplement 1.** CADA suppresses CD4 and CD8 receptor upregulation.

### CADA dose-dependently inhibits CD8^+^ T cell proliferation and cytotoxic T cell function

As CADA treatment resulted in lower expression of the CD8 receptor on CD8^+^ T cells, the effect of CADA on CD8^+^ T cell function was further examined. To this purpose, an MLR was performed with total PBMCs, purified CD4^+^ T cells or purified CD8^+^ T cells. Generally, the proliferation response of purified CD8^+^ T cells was much weaker for each donor in comparison to the proliferation response of purified CD4^+^ T cells (data not shown). As demonstrated in ***Figure 5A***, CADA dose-dependently suppressed the proliferation of purified CD4^+^ T cells, although to a lesser extent as compared to total PBMCs. Remarkably, CADA profoundly and dose-dependently inhibited the proliferation of purified CD8^+^ T cells, in a similar way as that of total PBMCs. In addition, the proliferation of purified CD8^+^ T cells by PHA or beads stimulation was clearly suppressed by CADA (***Figure 5B***). This indicates that the suppressive effect of CADA on lymphocyte activation is mostly affecting the CD8^+^ subpopulation and, thus, independent of CD4 expression.

**Figure 5.**
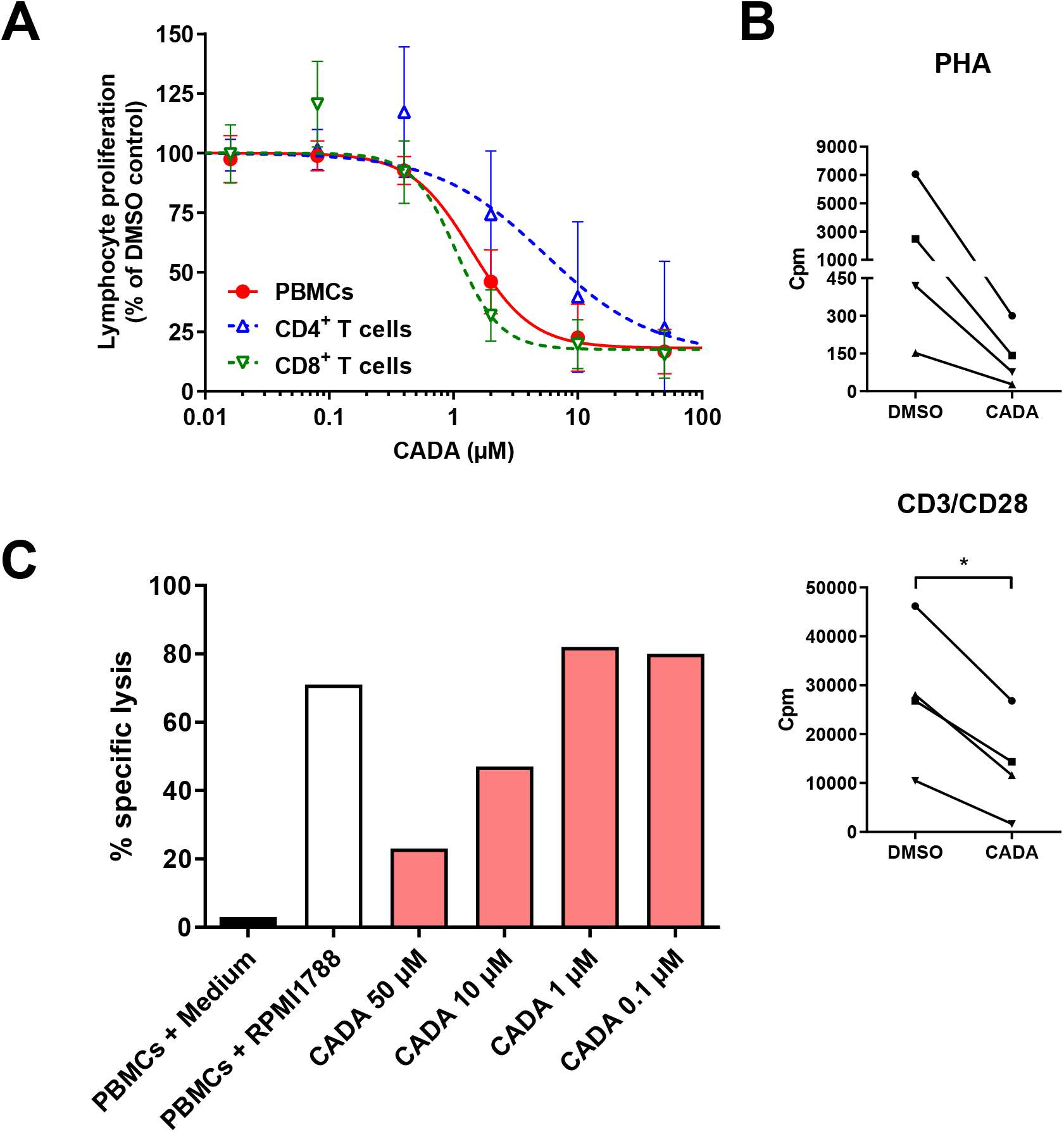
CADA dose-dependently inhibits CD8^+^ T cell proliferation and cytotoxic T cell function. **(A)** PBMCs (red), purified CD4^+^ T cells (blue) or purified CD8^+^ T cells (green) were co-cultured with mitomycin C inactivated RPMI1788 cells in the presence of different doses of CADA. At day 5, [^3^H]-thymidine was added and proliferation response was measured by detecting counts per minute (cpm) 18h later. Lymphocyte proliferation is given as percentage of the corresponding DMSO control (mean ± SD; n=6). **(B)** Purified CD8^+^ T cells were pre-incubated with CADA (10 μM) or DMSO during 3 days, after which they were activated by CD3/CD28 beads or PHA. At 24h post activation, [^3^H]-thymidine was added and proliferation response was measured by detecting cpm 20h later. Graphs show intra-donor treatment effect on the proliferation response (each donor is indicated separately). A paired t-test was performed to compare CADA to DMSO with *p<0.05. **(C)** PBMCs were cultured in medium alone (black) or were co-cultured with inactivated RPMI1788 cells in the absence (white) or presence (red) of increasing doses of CADA during 6 days. Next, PBMCs were co-cultured with ^51^Cr-loaded RPMI1788 cells for 4h, after which supernatant was collected and ^51^Cr release was quantified. To measure spontaneous and maximum release of ^51^Cr, medium or saponin was added to the ^51^Cr-loaded RPMI1788 cells, respectively. The mean percentage of specific lysis was calculated by using the following formula: % specific lysis = (experimental release – spontaneous release) / (maximum release – spontaneous release) × 100. Values of one experiment are shown.

Next, to evaluate the effect of CADA on the cytotoxic potential of CD8^+^ T cells, a cell-mediated lympholysis assay was performed. PBMCs cultured in medium without stimulator cells did not show notable cytotoxic activity (3% of specific lysis; black bar in ***Figure 5C***). However, when PBMCs were co-cultured with mitomycin C-inactivated RPMI1788 cells, cytotoxic activity increased considerably (71% of specific lysis; white bar in ***Figure 5C***). Interestingly, treatment with CADA reduced this cytotoxic response dose-dependently (77% inhibition of specific lysis with 50 μM of CADA, and 53% with 10 μM of CADA; red bars in ***Figure 5C***). At lower concentrations of CADA, cell-mediated lympholysis was no longer inhibited.

### CADA decreases CD25 upregulation and reduces intracellular pSTAT5 and CTPS1 levels in activated PBMCs

Expression of the late activation marker CD25 (also known as the low affinity IL-2 receptor α-chain) was determined on both CD4^+^ T cells and CD8^+^ T cells (***Figure 6A***). Without activation stimuli, very low levels of CD25 were measured, however, CD25 expression was strongly induced starting at 4h post PHA-activation, reaching a peak around day 2 to 3 (***Figure 6A***). Comparable data were obtained with CD3/CD28 beads activation (***Figure 6–figure supplement 1A***). Although CADA pre-incubation had no effect on basal CD25 levels (***Figure 6–figure supplement 1B***; d0), treatment of the cells with CADA inhibited CD25 upregulation in each T cell subset and in both activation models. As shown in ***Figure 6A*** (insert panels), CADA significantly suppressed CD25 expression at day 3 (p < 0.05; paired t-test). Though, at day 4 post activation the inhibitory effect of CADA was less distinct because CD25 expression already declined in most control samples, whereas it stabilized in CADA-treated cells (***Figure 6A***, ***Figure 6–figure supplement 1***). In accordance with cell surface expression of CD25, the level of soluble CD25 (sCD25) in the supernatant of stimulated cells was also reduced by CADA treatment (***Figure 6–figure supplement 2***), which was significant for the PHA-stimulated samples that were collected at day 4.

**Figure 6.**
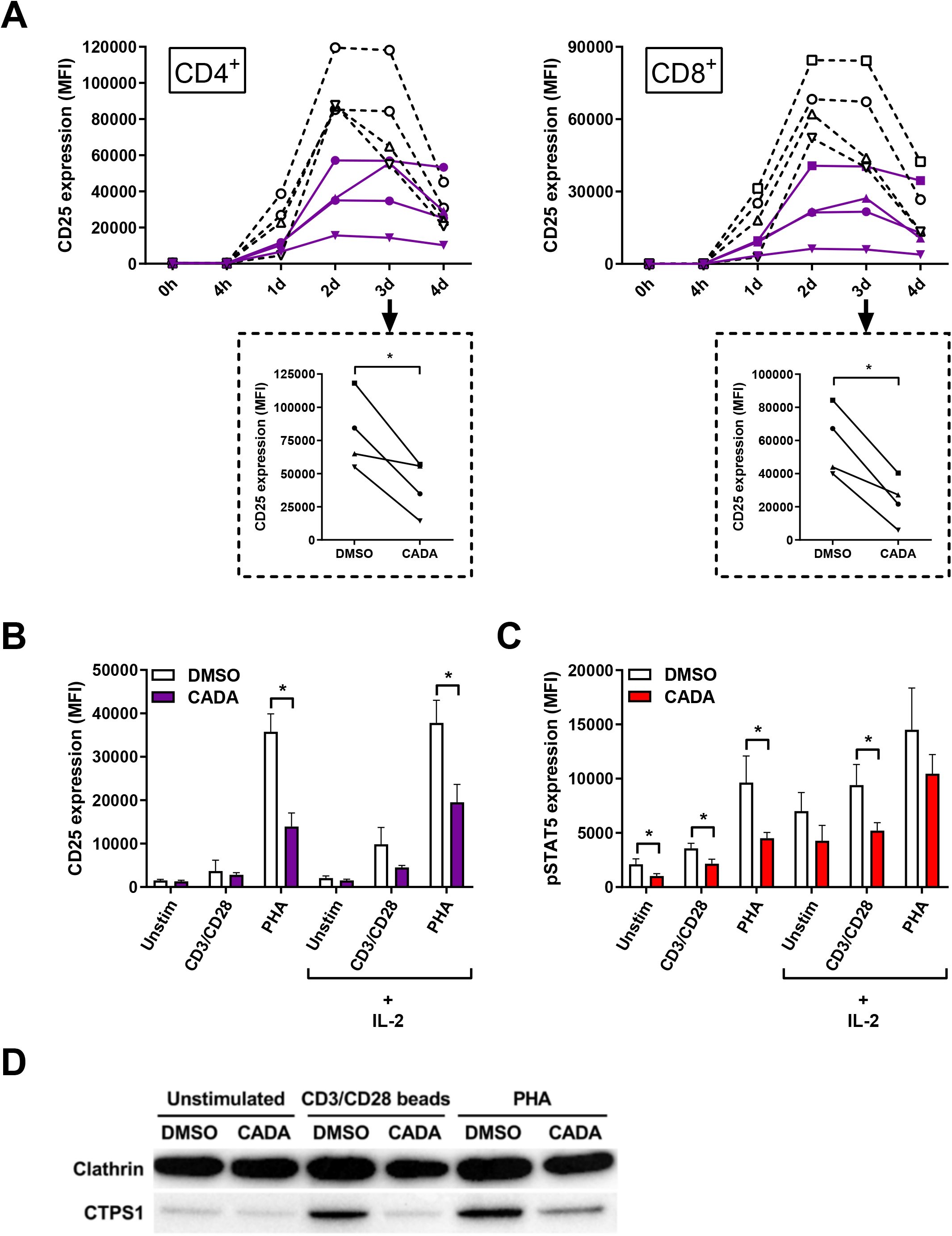
CADA decreases CD25 upregulation and reduces intracellular pSTAT5 and CTPS1 levels in activated PBMCs. **(A)** PBMCs were pre-incubated with CADA (10 μM) or DMSO for 3 days, after which they were activated with PHA. Cellular surface CD25 expression was measured on gated CD4^+^ (left panel) and CD8^+^ (right panel) T cells by flow cytometry just before activation (0h) and 4h, 1d, 2d, 3d or 4d post activation. Mean fluorescence intensity (MFI) of CD25 expression is shown for 4 donors of PBMCs (indicated separately) with DMSO-treated samples as a dotted line with open symbols and CADA-treated cells as a full purple line with solid symbols. Insert panels below each graph show intra-donor treatment effect on CD25 expression at day 3 post activation. A paired t-test was performed to compare CADA to DMSO with *p<0.05. **(B – D)** PBMCs were pre-incubated with CADA or DMSO during 3 days, after which they were left unstimulated or were activated with CD3/CD28 beads or PHA. **(B and C)** At day 2, half of the samples were boosted with IL-2. Cell surface CD25 receptor (B) and intracellular pSTAT5 (C) expression were simultaneously measured by flow cytometry. Mean fluorescence intensity (MFI) of CD25 and pSTAT5 is shown (mean ± SD; n=4). Multiple t-tests were performed to compare CADA (solid bars) to DMSO (open bars) for each condition with *p<0.05 and with Holm-Sidak method as correction for multiple comparison. **(D)** At day 2 post activation, cells were lysed and CTPS1 expression was detected by western blotting. Clathrin was used as protein loading control. **Figure supplement 1.** CADA inhibits CD25 upregulation after activation by CD3/CD28 beads. **Figure supplement 2.** CADA reduces the levels of soluble CD25 protein.

Transcription of CD25 is enhanced by IL-2 receptor signaling, including activation by phosphorylation of signal transducer and activator of transcription 5 (STAT5). Next, levels of intracellular pSTAT5 and cell surface CD25 were measured simultaneously in PBMCs that were left unstimulated or that were activated by CD3/CD28 beads and PHA. Half of the samples were given an extra boost with exogenous IL-2. As shown in ***Figure 6B***, most potent induction of CD25 expression in total PBMCs was obtained by PHA stimulation rather than by the use of CD3/CD28 beads. This CD25 upregulation, in the absence or presence of exogenous IL-2, was significantly suppressed by CADA (p = 0.001 and 0.007, respectively; t-test). Activation with CD3/CD28 beads, in combination with exogenous IL-2 also resulted in detectable levels of CD25 (***Figure 6B***). Intracellular pSTAT5 levels were clearly elevated after activation, with the largest increase for the PHA-stimulated samples (***Figure 6C***). By adding exogenous IL-2 a general increase in pSTAT5 was seen in all tested conditions. Interestingly, CADA clearly reduced the levels of pSTAT5 (as compared to the corresponding DMSO control), which reached significance for the samples without IL-2 boost (***Figure 6C***, red bars). In addition, the expression level of cytidine triphosphate synthase 1 (CTPS1) – an important immune checkpoint in T cell responses – was determined as its transcription is induced by activated STAT5. CTPS1 is highly upregulated after stimulation and it has been reported to be crucial for proliferation of T and B cells after activation (Martin et al., 2014). As shown in ***Figure 6D***, in unstimulated cells low basal levels of CTPS1 were detected by means of western blot, while enhanced expression was observed after activation by CD3/CD28 beads and PHA. Importantly, CADA clearly attenuated this activation-induced CTPS1 upregulation (***Figure 6D***).

### CADA inhibits cytokine release by activated PBMCs and suppresses the upregulation of co-stimulatory molecules

Our first set of data indicated that CADA attenuates the general activation of T lymphocytes. To explore the immunosuppressive effect of CADA in more detail, we next analyzed the impact of CADA on the cytokine release by the proliferating lymphocytes. Supernatant was taken from PBMCs either stimulated by mitomycin C inactivated RPMI1788 cells (MLR), CD3/CD28 beads or PHA and analyzed for three representative Th1 cytokines. As summarized in ***Figure 7A***, CADA generally suppressed the level of IL-2, IFN-γ and TNF-α in the three activation models, which reached significance for the cytokines detected in the MLR samples. TNF-α was significantly reduced by CADA treatment in all three models (p<0.05; t-test).

**Figure 7.**
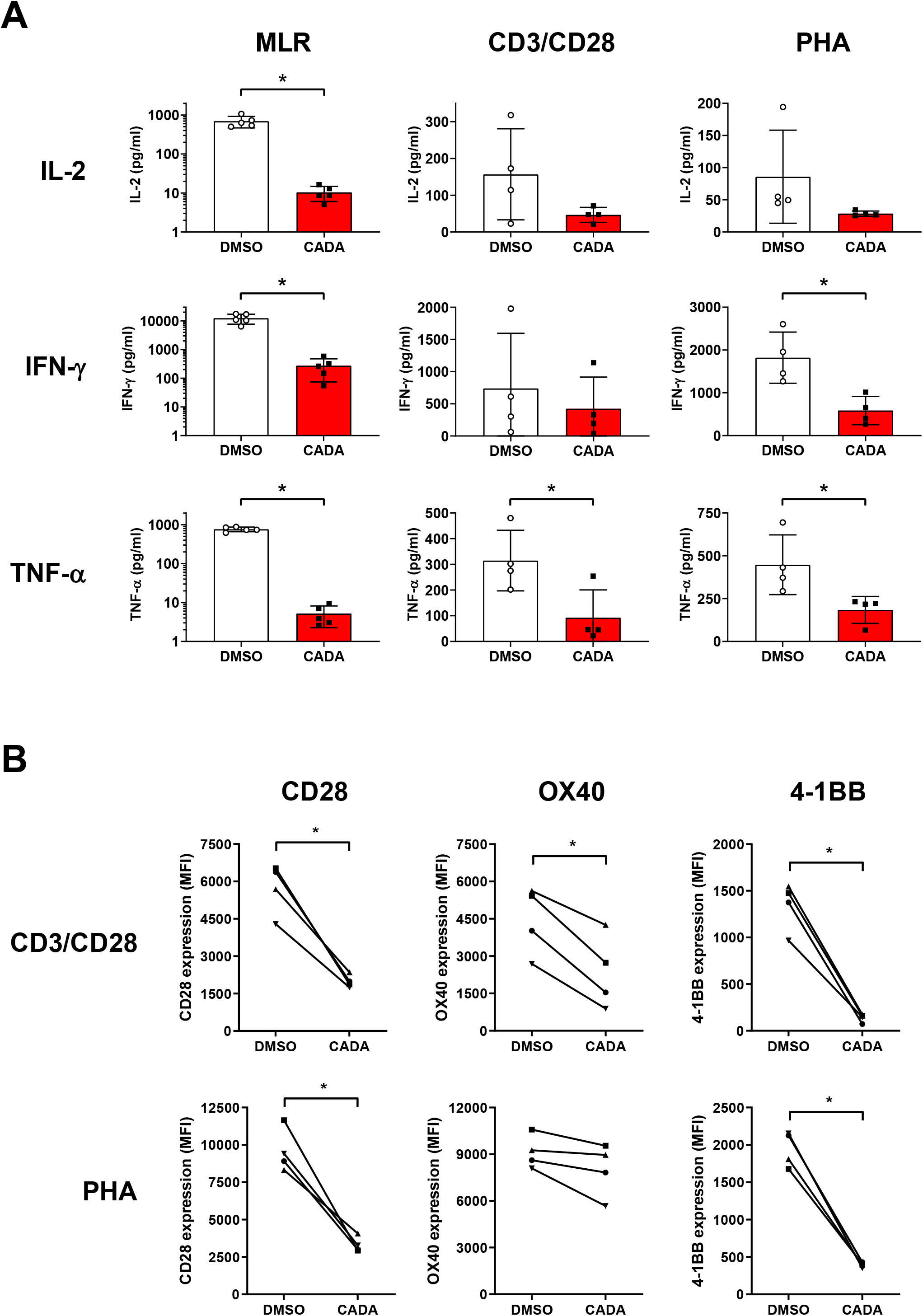
CADA inhibits cytokine release by activated PBMCs and suppresses the upregulation of co-stimulatory molecules. **(A)** PBMCs were stimulated with mitomycin C inactivated RPMI1788 cells (MLR), CD3/CD28 beads or PHA and exposed to CADA (10 μM). Supernatants were collected on day 5 (MLR; n=5) or day 3 (beads and PHA; n=4) post stimulation and cytokine levels were determined by Bio-Plex assay. Bars represent mean ± SD, with individual values shown as open (DMSO) or solid (CADA) symbols. Note that cytokine levels in the MLR samples are plotted on a logarithmic scale. Welch’s corrected t-tests were performed to compare CADA to DMSO with *p<0.05. **(B)** PBMCs were pre-incubated with CADA (10 μM) or DMSO during 3 days, after which they were activated by CD3/CD28 beads or PHA. Cell surface CD28 expression was measured on gated CD4^+^ T cells by flow cytometry on d3 post activation. Cell surface expression of OX40 and 4-1BB was measured on total PBMCs by flow cytometry on d2 post activation. Panels represent intra-donor treatment effect of CADA on receptor expression for 4 donors of PBMCs (indicated separately). Paired t-tests were performed to compare CADA to DMSO with *p<0.05. **Figure supplement 1.** CADA reduces the upregulation of CD28 on activated lymphocytes.

In addition to the cytokine response of lymphocytes, we evaluated the expression level of CD28, a key co-stimulatory receptor in T cell activation. As depicted in ***Figure 7–figure supplement 1***, cell surface CD28 expression levels started to rise at day 2 post activation. Treatment with CADA resulted in a significant reduction in CD28 levels of CD4^+^ and CD8^+^ T cells, both after CD3/CD28 and PHA stimulation (***Figure 7B, Figure 7–figure supplement 1***). By day 3 post activation, CD28 expression levels generally increased also in the CADA-exposed samples (***Figure 7–figure supplement 1***), indicating that CADA-treatment resulted in a delayed upregulation of CD28 rather than a complete and sustained suppression of this co-receptor.

Cell surface levels of the human co-stimulatory receptors tumor necrosis factor receptor superfamily [TNFRSF] member 4 (TNFRSF4), also named OX40 or CD134, and 4-1BB (also named CD137 or TNFRSF9) were assessed after activation by CD3/CD28 beads or PHA. OX40 is transiently expressed after antigen recognition primarily on activated CD4^+^ T cells found preferentially at the site of inflammation (Lane, 2000; Weinberg, 2002). Expression of 4-1BB is highly induced in CD8^+^ T and NK lymphocytes upon activation via CD3-TCR engagement. It exerts regulatory effects on T cells mediating activation and persistence of CD8^+^ T lymphocytes (Kwon et al., 2000; Shuford et al., 1997; Vinay et al., 1998). As shown in ***Figure 7B***, activation of the control cells evoked a strong but variable upregulation of OX40, with higher elevated levels after stimulation with PHA as compared to CD3/CD28 beads activation. The suppressive effect of CADA on OX40 upregulation was rather weak in PHA-stimulated cells (13 ± 12% reduction in MFI), though it was more pronounced (51 ± 19% reduction in MFI) and reached statistical significance in the case of CD3/CD28 beads activation (p = 0.0064; paired t-test). The most striking effect was observed for 4-1BB expression. In both activation models, an uniform increase in 4-1BB expression was measured in the DMSO control samples of the four different donors (***Figure 7B***). In sharp contrast, CADA nearly completely blocked the upregulation of 4-1BB in all samples (89 ± 4% and 79 ± 3% reduction in MFI for CD3/CD8 and PHA, respectively), which was highly significant (p = 0.0025 and 0.0011, respectively; paired t-test).

### CADA dose-dependently and reversibly suppresses the cellular expression of 4-1BB

Further analysis of 4-1BB kinetics indicated that the transient expression of 4-1BB in CD8^+^ T cells starts as early as 12h post stimulation and lasts for approximately 36h, whereas its expression in CD4^+^ T cells peaks around 48h post stimulation (***Figure 8***). Importantly, CADA completely abrogated the 4-1BB upregulation in both CD8^+^ and CD4^+^ T cells (***Figure 8***, red curves). These data suggest that CADA might have a direct inhibitory effect on the receptor biogenesis of 4-1BB, similar to that of CD4. To address this, we cloned 4-1BB in a vector to express the receptor fused to turbo green fluorescent protein (tGFP) in a P2A-RFP context (***Figure 9A***), as described previously (Van Puyenbroeck et al., 2020). As a positive control, hCD4 was included. The same reporter vector was also used to express other co-stimulatory receptors from the same genetic background. Protein expression was determined by tGFP fluorescence, while the amount of cytosolic RFP served as a control for transfection and expression efficiency. As shown in ***Figure 9B***, CADA dose-dependently inhibited 4-1BB expression in transfected HEK293T cells. This direct down-modulatory effect of CADA on 4-1BB was almost complete and similar to its effect on hCD4 (IC_50_ of 0.24 μM and 0.30 μM, respectively), demonstrating that 4-B11 is a valuable substrate of CADA (***Figure 9B***). The down-modulating effect of CADA on 4-1BB is reversible in nature, as evidenced by the re-expression of 4-1BB after wash-out of CADA (***Figure 9C***), an effect that is observed for hCD4 as well (***Figure 9–figure supplement 1***) as reported earlier (Vermeire et al., 2014; Vermeire et al., 2007). As summarized in ***Figure 9D***, in addition to the potent inhibition of hCD4 and 4-1BB expression, CADA also partially reduced cellular levels of other co-stimulatory receptors in transfected cells. Whereas the level of CD8 and OX40 in CADA treated cells was reduced by approximately 40% the effect of CADA on the expression of CD25 and CD69 was only minor. A reduction of 60% was measured in the expression of CD28 in CADA exposed cells (***Figure 9D, Figure 9–figure supplement 2***).

**Figure 8.**
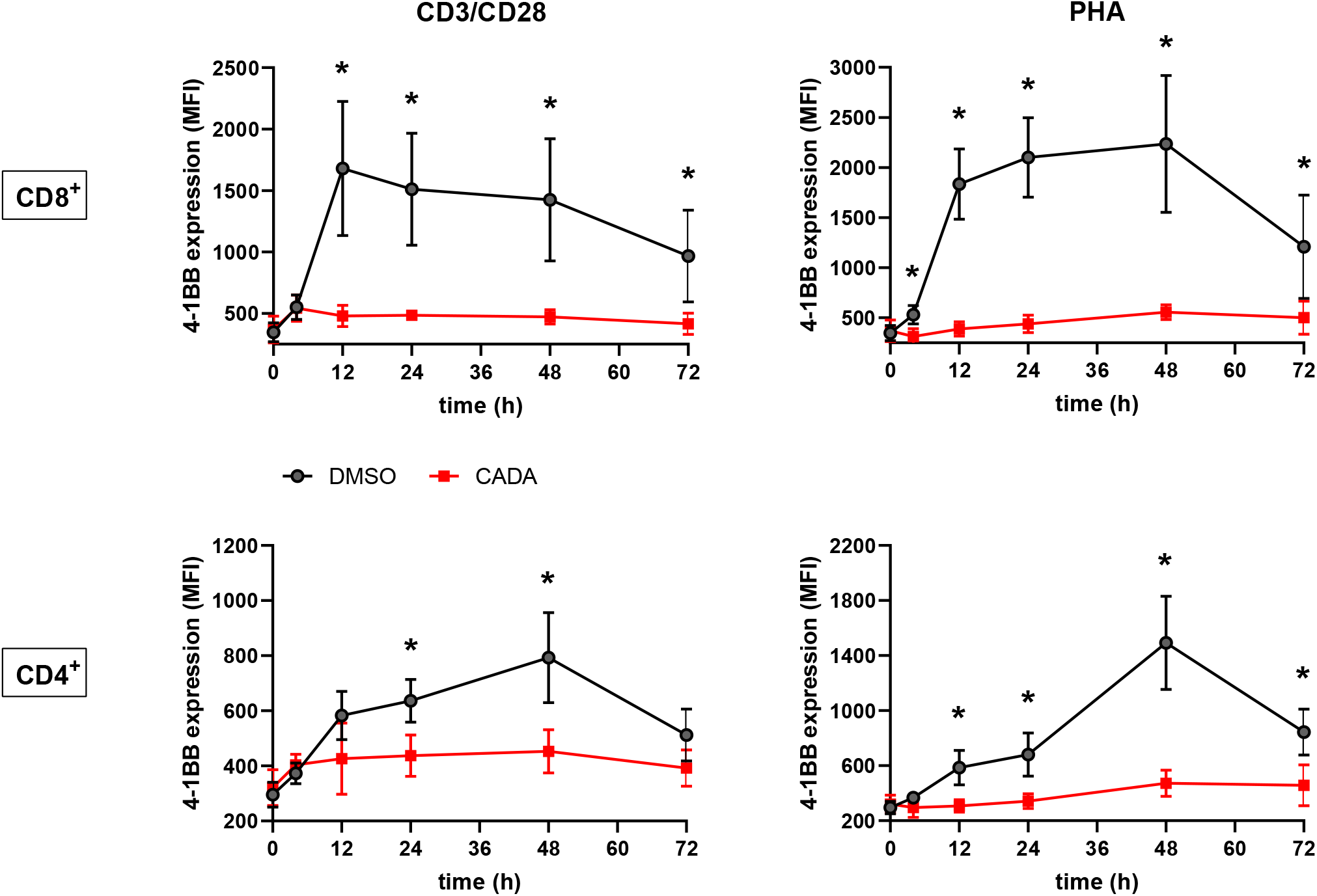
CADA completely suppresses the upregulation of 4-1BB. PBMCs were pre-incubated with CADA (10 μM) or DMSO during 3 days, after which they were activated by CD3/CD28 beads or PHA. Cell surface 4-1BB expression was measured on gated CD4^+^ and CD8^+^ T cells by flow cytometry on the indicated time points post activation. The average MFI of 6 donors of PBMCs is shown (mean ± SD). Multiple t-tests were performed to compare CADA to DMSO for each condition with *p<0.05 and with Holm-Sidak method as correction for multiple comparison.

**Figure 9.**
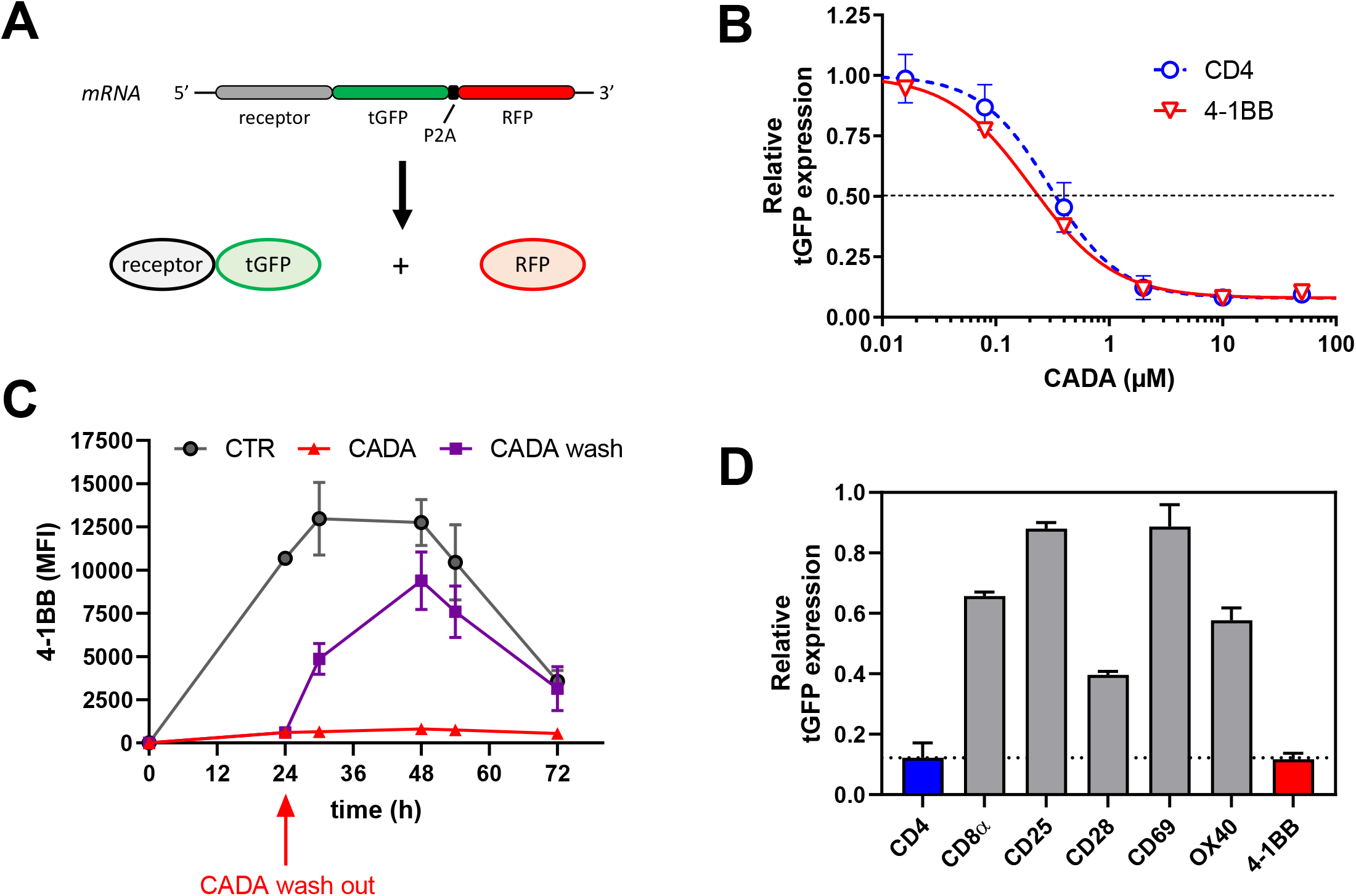
CADA dose-dependently and reversibly suppresses the cellular expression of 4-1BB. **(A)** Schematic representation of the expected mRNA and protein products of the tGFP-2A-RFP construct. **(B-D)** HEK293T cells were transiently transfected with the different constructs. CADA was added 6h post transfection and cellular expression of each receptor was determined by measuring tGFP levels by flow cytometry. **(B)** Four parameter dose-response curves for CADA of human CD4tGFP-2A-RFP and human 4-1BBtGFP-2A-RFP. Cells were collected 24h post transfection and tGFP was measured by flow cytometry. Receptor levels in CADA-treated samples are normalized to the corresponding DMSO control. Values are mean ± SD; n ≥ 3. **(C)** Cells were transfected with 4-1BBtGFP-2A-RFP and given DMSO (CTR) or treated with CADA for 72h. In parallel, CADA-treatment was terminated after 24h (CADA wash). These cells were washed profoundly and given control medium for the duration of the experiment. At the indicated time points, cells were collected and tGFP was measured by flow cytometry. The average MFI of tGFP is shown (mean ± SD; n=2). Of note is that the SD of the CADA samples (red curve) is too small to be visible on the graph. **(D)** Cells were collected 24h post transfection and tGFP was measured by flow cytometry. Protein levels in CADA-treated samples are shown, normalized to the corresponding DMSO control (set as 1.00). Bars are mean ± SD; n ≥ 3. **Figure supplement 1.** CADA reversibly down-modulates hCD4. **Figure supplement 2.** CADA differentially affects the protein expression levels of co-stimulatory receptors in transfected cells.

### CADA inhibits 4-1BB protein biogenesis is a signal peptide-dependent way by blocking the co-translational translocation of 4-1BB into the endoplasmic reticulum

Finally, to explore the molecular mechanism by which CADA inhibits 4-1BB protein expression, we addressed if the cleavable signal peptide (SP) of the 4-1BB pre-protein is the susceptible region for CADA activity, similar to what we have described for hCD4 (Van Puyenbroeck et al., 2020; Vermeire et al., 2014). Thus, constructs were generated as depicted in ***Figure 10A***. Briefly, starting from the CADA-resistant mouse CD4 (mCD4) protein sequence, we exchanged the N-terminal region containing the SP and the first 7 amino acids of the mature protein of mCD4 with that of hCD4 or 4-1BB, respectively. As previously demonstrated (Vermeire et al., 2014), CADA did not affect the expression of wild-type mouse CD4 when transfected in HEK293T cells (***Figure 10B***). Expectedly, mCD4 could be fully sensitized to CADA by substituting the mCD4 SP and the first 7 amino acids of the mature mCD4 protein by the human sequence (hmCD4 construct; ***Figure 10B***), confirming that CADA-sensitivity depends on the presence of a hCD4 SP. Interestingly, expression of mouse CD4 could also be dose-dependently down-modulated by CADA when mCD4 contained the 4-1BB SP and 7 AA of the mature 4-1BB protein. In fact, 4-1BBmCD4 was slightly more affected by CADA as compared to the hmCD4 chimaera, as evidenced by the IC_50_ values for receptor down-modulation (0.38 and 0.84 μM, respectively) (***Figure 10B***).

**Figure 10.**
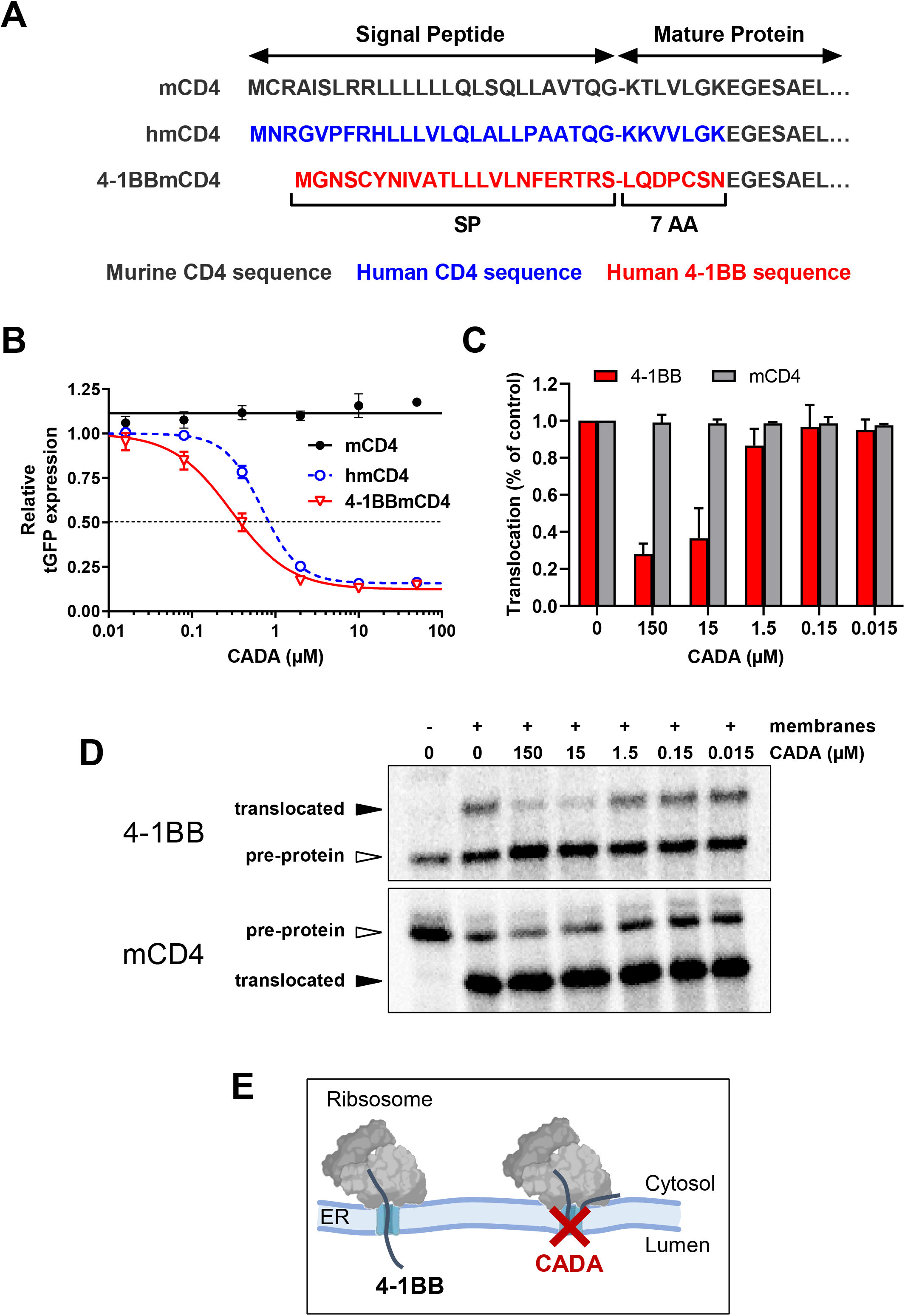
CADA inhibits 4-1BB protein biogenesis is a signal peptide-dependent way by blocking the co-translational translocation of 4-1BB into the endoplasmic reticulum. **(A)** Schematic representation of the constructs used. In the hmCD4 construct, the signal peptide (SP) and the first 7 amino acids of the mature protein are of human CD4 (indicated in blue), whereas in the 4-1BBmCD4 construct the SP and the 7 AA of mature 4-1BB (indicated in red) are fused to mouse CD4. During pre-protein biogenesis, the SP is cleaved off from the mature protein. The constructs express the mature protein of mouse CD4 that is C-terminally fused to tGFP as shown in Figure 9A. **(B)** HEK293T cells were plated and transfected with the mCD4 (black; n=3), hmCD4 (blue; n=3) and 4-1BBmCD4 (red; n=4) constructs. CADA was added 6h post transfection and expression of tGFP was measured by flow cytometry 24h post transfection. The tGFP expression is given as percentage of DMSO control (mean ± SD). IC_50_ values are 0.84 μM, 0.38 μM and >50 μM for hmCD4, 4-1BBmCD4 and mCD4, respectively. **(C and D)** *In vitro* translation and translocation of 4-1BB and mCD4 in a radiolabeled cell-free rabbit reticulocyte lysate system. **(C)** Graph shows the calculated translocation efficiencies. Signal intensities of the pre-protein and translocated protein fraction were used to calculate the translocation efficiency, i.e., translocated fraction/(pre-protein + translocated fraction). Bars show mean ± SD; n=2. **(D)** Representative autoradiogram of the *in vitro* translated and translocated wild-type 4-1BB and mCD4 proteins. For mCD4 a truncated form of 250 residues was used without glycosylation sites and transmembrane region. In the presence of membranes, the pre-protein (open arrowhead) of mCD4 is translocated into the ER lumen and the SP is cleaved off, resulting in a faster migrating mature protein (black arrowhead). For wild-type 4-1BB, the SP is cleaved off but the protein is also glycosylated, resulting in a slower migrating mature protein (black arrowhead). **(E)** Cartoon showing CADA inhibiting the co-translational translocation of 4-1BB protein across the ER membrane. **Figure supplement 1.** Cell free *in vitro* translation/translocation assay to study the co-translational translocation of proteins.

Signal peptides are critical targeting sequences for secretory and type I integral membrane proteins to guide these proteins to the secretory pathway (von Heijne, 1985; Wickner et al., 2005). They are involved in the correct targeting of translating ribosomes to the endoplasmic reticulum (ER) membrane, and the subsequent selective translocation of secretory and type I integral membrane proteins across the Sec61 translocon channel in the ER membrane (Hegde et al., 2008; Rapoport, 2007). By the use of a cell free *in vitro* translation/translocation assay (Vermeire et al., 2015), we next evaluated the impact of CADA specifically on the translocation step of 4-1BB (***Figure 10–figure supplement 1***). Briefly, transcripts of full length 4-1BB were translated *in vitro* into a pre-protein of approximately 30 kDa, containing its SP (***Figure 10D***, top panel, first lane). By adding microsomal membranes, representing the ER, combined translation and translocation can occur, resulting in SP-cleaved proteins that are further glycosylated in the ER lumen by the oligosaccharyltransferase (OST) complex (***Figure 10– figure supplement 1A***). As shown in ***Figure 10D***, wild-type 4-1BB is efficiently translocated into the lumen of the microsomal membranes, as evidenced by the higher molecular weight band on the gel representing the translocated (thus, glycosylated) 4-1BB species. However, addition of CADA to this translocation mixture strongly reduced the fraction of translocated protein, demonstrating that CADA specifically inhibits the protein translocation step of 4-1BB (***Figure 10C***). In contrast, CADA had no effect on the translocation of wild-type truncated mCD4 (without glycosylation sites), as evidenced by the equal amount of faster migrating SP-cleaved species (***Figure 10C*** and ***Figure 10D***, bottom panel). These data demonstrate that CADA specifically inhibits the co-translational translocation of 4-1BB across the ER membrane in a signal peptide-dependent manner (***Figure 10E***).

## DISCUSSION

This study aimed at evaluating the immunosuppressive potential of CADA, a small molecule that blocks hCD4 protein biosynthesis in a SP-dependent way and thereby reduces cell surface hCD4 expression to low basal level. Here, we demonstrated a consistent dose-dependent inhibitory effect of CADA on lymphocyte proliferation in a MLR setting. The inhibition of lymphocyte proliferation by CADA was milder than by the currently used anti-proliferative immunosuppressive agent MMF. Although less potent, CADA has the major advantage that it exerted no cellular toxicity and it was barely cytostatic *in vitro*, both promising beneficial characteristics of an immunosuppressive drug. In addition, the biological effect of CADA is reversible as evidenced by the quick re-expression of the targeted receptors when treatment was terminated. CADA had little suppressive effect on superantigen-induced activation of T cells. This can be explained by the unique binding of superantigens, which occurs outside the normal peptide-binding groove and thus without intracellular processing (Marrack et al., 1990). Interactions between superantigen and TCR or MHC are most likely of sufficiently high affinity to obviate the contribution of the CD4 receptor in this activation process (Killeen et al., 1993), explaining the lack of a significant suppressive effect of CADA which was expected to be mainly CD4-based. With additional data generated in two different *in vitro* T cell activation models (i.e., CD3/CD28 beads and PHA), we confirmed that CADA significantly inhibits the proliferation response of stimulated lymphocytes. Although the activation signals in T cells in these models are weakened but not completely blocked by CADA, this partial and temporal suppressive effect of CADA is certainly meaningful. Notably, the supra physiological stimulation of T cells with both CD3/CD28 beads and PHA is a condition that is never achieved in a normal *in vivo* setting where only a small subset of T cells is selectively triggered.

When comparing the active dose ranges of CADA with Clenoliximab in the MLR, we concluded that CADA was more potent than Clenoliximab at down-modulating hCD4 expression and at inhibiting lymphocyte proliferation. Clenoliximab is a nondepleting anti-CD4 monoclonal antibody that directly targets the hCD4 receptor (Hepburn et al., 2003). This antibody reached phase II clinical trial for the treatment of rheumatoid arthritis (Mould et al., 1999). The concentrations of Clenoliximab used in our study were considered adequate to obtain maximum activity, as previously an IC_50_ of 14.6 ng/ml Clenoliximab was reported in the MLR (Reddy et al., 2000). However, in our hands Clenoliximab exerted only a partial immunosuppressive effect, but this may be due to different assay characteristics (Reddy *et al.* used a three-way MLR, whereas we performed a one-way MLR). Either way, the data presented here indicate that the immunosuppressive capacity of CADA in the MLR exceeded that of Clenoliximab. Remarkably, at concentrations of 50, 10 and 2 μM of CADA similar down-modulation of hCD4 was elicited, whereas the inhibitory effect of CADA on lymphocyte proliferation still increased with higher concentrations (***Figure 2***). This suggested that besides reduction in CD4 expression, other factors may be at play in the total immunosuppressive effect of CADA.

Interestingly, CADA inhibited the proliferation of purified CD8^+^ T cells to the same extent in the absence of other immune cell types as compared to the proliferation of total PBMCs in the MLR. In addition, the proliferation of purified CD8^+^ T cells by stimulation with CD3/CD28 beads or PHA was also clearly suppressed by CADA treatment. Moreover, CADA inhibited cytotoxic cell activity in a cell-mediated lympholysis assay. These data demonstrate a direct inhibitory effect of CADA on CD8^+^ T cell proliferation and function, independently of CD4 receptor expression. This effect cannot solely be attributed to reduced CD8 receptor levels measured in the cytotoxic T cells, as CADA suppressed CD8 levels only partially. Similar to the function of CD4 on CD4^+^ T cells, the CD8 receptor enhances the sensitivity of CD8^+^ T cells to antigens and is required for the formation of a stable complex between major histocompatibility complex class I and the T cell receptor (Xiao et al., 2007). However, the nearly complete inhibition of 4-1BB upregulation in CD8^+^ cells is most likely one of the main raisons for the strong non-CD4 dependent immunosuppression of CADA in the CD8^+^ T cell population. Indeed, a clear role of 4-1BB in augmenting T cell cytotoxicity and CD8^+^ T cell survival has been reported in literature (Kwon et al., 2000; Shuford et al., 1997; Vinay et al., 1998). The surface glycoprotein 4-1BB is a member of the TNFR family whose expression is highly induced in CD8^+^ T and NK lymphocytes upon activation via CD3-TCR engagement. It functions as an inducible co-stimulatory molecule that can exert regulatory effects on T cells mediating activation and persistence of cytotoxic T lymphocytes independently of CD28 stimulation (Cannons et al., 2001; Kwon et al., 1989; Lin et al., 2008; Melero et al., 1998; Shuford et al., 1997; Wang et al., 2003). The finding that 4-1BB-mediated co-stimulation is critical for CD8^+^ T cell responses is further underlined in 4-1BB deficient mice in which decreased IFN-γ production and cytolytic CD8^+^ T cell effector function were observed (Kwon et al., 2002). In addition, 4-1BB deficiency in patients resulted in defective CD8^+^ T cell activation and cytotoxicity against virus-infected B cells (Alosaimi et al., 2019).

From our molecular biology data, we concluded that 4-1BB is an additional substrate of CADA in the context of co-translational protein translocation inhibition across the ER membrane during early protein biogenesis. This process involves the SP of the pre-protein for inserting into the translocon channel of the ER and correct routing along the secretory pathway (Hegde et al., 2008; Rapoport, 2007; von Heijne, 1985; Wickner et al., 2005). Although originally assumed that hCD4 was the sole target of CADA (Vermeire et al., 2014), a recent proteomic study indicated sortilin as a secondary substrate of CADA but with reduced sensitivity to the drug (Van Puyenbroeck et al., 2017). In an additional proteomics analysis of SUP-T1 cells (which is still ongoing), only a few hits out of more than 3000 quantified integral membrane proteins could be identified as susceptible to CADA but all with weaker sensitivity as compared to hCD4. Also in our current study it is clear that CADA has not a general inhibitory effect on protein translocation of the total integral membrane fraction as evidenced by CD25 and CD69 whose expression in transfected cells was unaffected by CADA. From our comparative analysis in transfected cells we can now conclude that 4-1BB is the most sensitive substrate of CADA identified so far, making it an ideal target for further mechanistic studies. By comparison with hCD4 we hope to get a better understanding of how a small molecule can exert such a high substrate selectivity for ER translocation inhibition in order to ultimately design novel ER translocation inhibitors for therapeutic use.

The upregulation of several immunologically relevant receptors after T cell stimulation was shown to be suppressed by CADA. To distinguish between reduced expression level because of a general immunosuppression by CADA and a direct inhibition of protein translocation and subsequent receptor expression, we evaluated the expression efficiency of each receptor independently in transfected cells. Unaffected by CADA directly, the expression of late activation marker CD25 – also known as IL-2 receptor α-chain – was significantly reduced and somewhat delayed by CADA after activation with CD3/CD28 beads and PHA. Thus, the CD25 expression level in CADA-exposed activated T cells is a relevant measurement of the degree of actual T cell activation. This can also explain the higher variation in CD25 expression level between the different CADA-treated donors (***Figure 6A***) as compared to hCD4 (***Figure 4A***). Expectedly, we also observed a decreased amount of sCD25 in the supernatant of activated lymphocytes. sCD25 is a sensitive marker for activation of the immune system and it can also be used as a potential marker for subclinical macrophage activation syndrome in patients with active systemic onset juvenile idiopathic arthritis (Reddy et al., 2014). CD25 expression is massively upregulated after T cell activation involving T cell receptor and IL-2 receptor signaling pathways (Shatrova et al., 2016). In the IL-2 receptor signaling pathway, activation of STAT5 by phosphorylation is crucial to enhance CD25 expression. Furthermore, cytidine triphosphate synthase 1 (CTPS1) transcription is induced by activated STAT5, and as an enzyme in the *de novo* synthesis of cytidine triphosphate, CTPS1 is crucial for proliferation of activated T and B cells (Martin et al., 2014). Its expression is rapidly and strongly upregulated following T cell activation. CTPS1 plays a predominant role in selected immune cell populations – e.g. CTPS1-deficient patients present with a life-threatening immunodeficiency – making CTPS1 an interesting target for the development of highly selective immunomodulatory drugs. CADA-treatment not only resulted in reduced CD25 and pSTAT5 levels, but also in reduced down-stream CTPS1 expression. Together with the suppressed release of pro-inflammatory cytokines, these data support our conclusion of CADA’s immunosuppressive potential.

A major co-stimulatory receptor in T cell activation is CD28. Treatment of the cells with CADA clearly inhibited the upregulation of CD28. This was partially the result of direct CADA-inhibition on CD28 protein expression. The inhibitory effect of CADA on CD28 was not complete, as evident from the residual expression (about 50%) in activated cells, but certainly meaningful.

Blocking CD28 has been shown to be successful in inhibiting unwanted T cell responses and the use of CADA would circumvent the risk of generating an agonistic signal, as is potentially the case for anti-CD28 monoclonal antibodies (Beyersdorf et al., 2015). Also, as 4-1BB is able to replace CD28 in stimulating high-level IL-2 production by resting T cells in the absence of CD28 (Chu et al., 1997; DeBenedette et al., 1997; Saoulli et al., 1998), the combined inhibition of signaling through CD28 and 4-1BB by CADA provides an interesting additional effect. Both co-stimulatory factors have sequentially differential roles in the stages of immune response with CD28 involved in the induction stage and 4-1BB in perpetuating the immune response providing a survival signal for T cells (Hurtado et al., 1997; Kwon et al., 2000; Takahashi et al., 1999).

In this study, 4-1BB has been discovered as a new target of CADA. Recently, the role of 4-1BB agonistic signaling in cancer immunotherapy has received great attention: the effect of 4-1BB stimulation by means of agonistic monoclonal anti-4-1BB antibodies on cytolytic T-cell responses has been used to increase the potency of vaccines against cancers (Bartkowiak et al., 2015; Chester et al., 2016; Oda et al., 2020). Therapeutic use of CADA would imply depletion of 4-1BB in order to attenuate cytotoxic T cell activity. In this context, blockade of 4-1BB has been shown to significantly impair the priming of alloantigen-specific CD8^+^ T cells and to increase allograft survival after transplantation (Cho et al., 2004; Wang et al., 2003), thus, suggesting a valuable application for CADA as new immunosuppressive drug in the field of e.g., organ transplantation. Furthermore, in the more general context of inflammatory diseases with a role of the adaptive immunity, general immunosuppression by CADA might be relevant to control, for instance, cytokine storm in hemophagocytic lymphohistiocytosis (HLH), severe cytokine release syndrome (CRS) in CAR T cell treatment, or even auto-immune diseases. As mainly human targets have been identified for CADA and resistance has been observed for e.g., murine CD4, humanized *in vivo* animal models are needed to fully evaluate CADA’s potential in these fields.

In conclusion (***Figure 11***), we showed here that the ER translocation inhibitor CADA exerted a profound and consistent *in vitro* immunosuppressive effect in the MLR and after activation with CD3/CD28 beads or PHA. This immunosuppressive effect of CADA involves both CD4^+^ and CD8^+^ T cells, but is most prominent in the CD8^+^ T cell subpopulation where it inhibits cell-mediated lympholysis. Next to the full suppression of CD4 and 4-1BB receptor upregulation, the combined effect of CADA on additional co-stimulatory factors such as CD28, OX40 and CD8 characterize the total immunosuppressive potential of CADA. Taken together, our data justify future *in vivo* exploration of this compound to evaluate its potential use to repress undesired immune activation.

**Figure 11.**
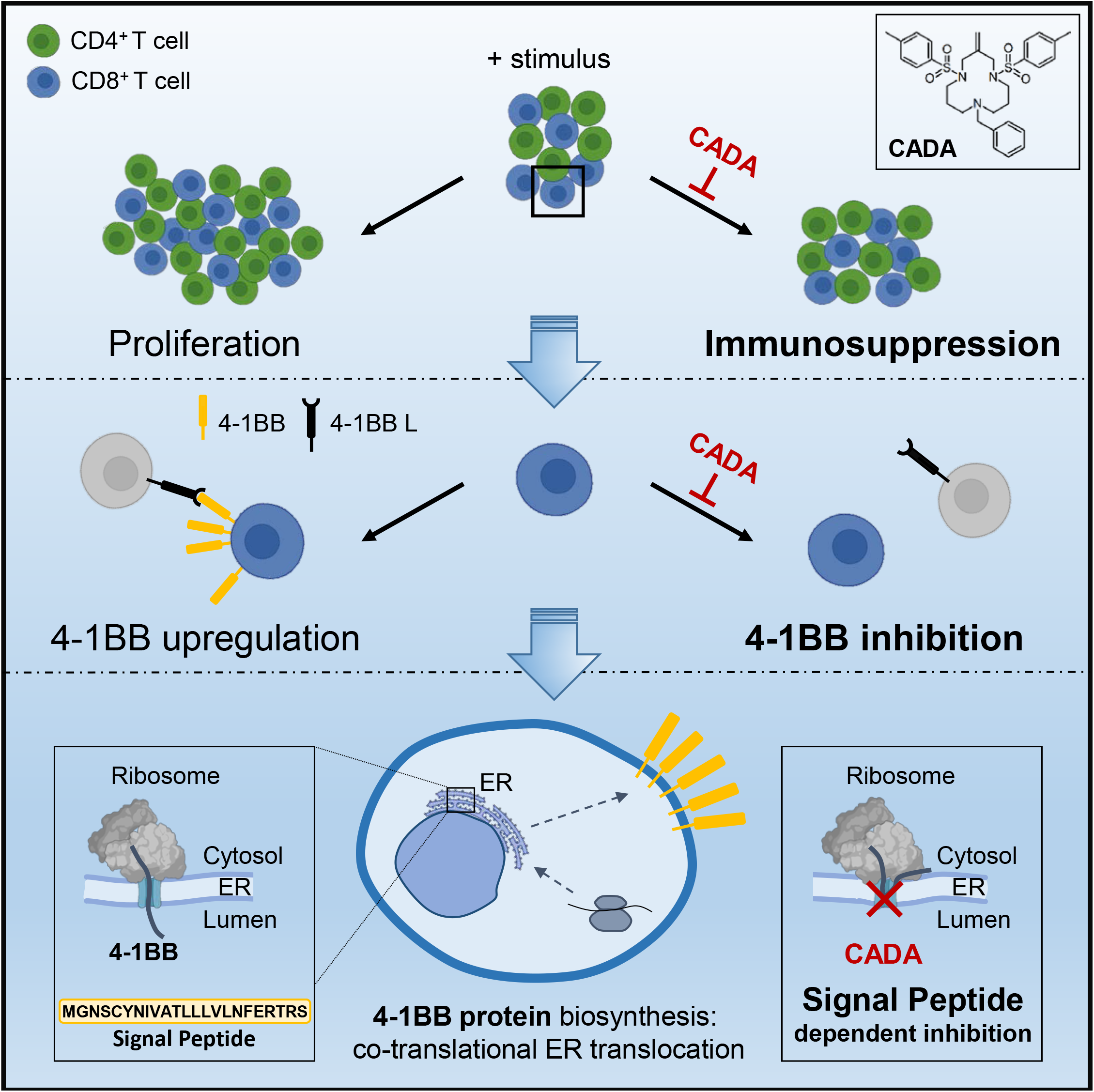
Mode of action of CADA. CADA has immunosuppressive activity mainly on CD8^+^ T cells by inhibition of 4-1BB protein biogenesis is a signal peptide-dependent way.

## MATERIAL AND METHODS

### Compounds and antibodies

CADA was a gift from Dr. Thomas W. Bell (University of Nevada, Reno). It was synthesized as described previously (Bell et al., 2006). Mycophenolate mofetil (MMF) was obtained from Sigma-Aldrich. Both compounds were dissolved in dimethyl sulfoxide (DMSO) to obtain a 10 mM stock solution for use in cell culture. The anti-CD4 monoclonal antibody Clenoliximab (chimeric macaque/human IgG4 antibody) was purchased from Absolute Antibody. Flow cytometry antibodies were purchased from (i) eBioscience (Thermo Fisher Scientific): APC-labeled anti-mouse CD4 (clone GK1.5) and APC-labeled anti-human phospho-STAT5 (Tyr694) (clone SRBCZX); (ii) BioLegend: PE-labeled anti-human CD4 (clone SK3), PE-labeled anti-human CD4 (clone OKT4), APC-labeled anti-human CD4 (clone SK3) and PE-labeled anti-human CD69 (clone FN50); (iii) BD Biosciences: BV510-labeled anti-human CD8 (clone SK1), PE-labeled anti-human CD25 (clone 2A3), FITC-labeled anti-human CD25 (clone 2A3), PE-labeled anti-human CD28 (clone CD28.2), BV421-labeled anti-human GITR (clone V27-580), PE-labeled anti-human OX40 (clone ACT35), PE-labeled anti-human 4-1BB (clone 4B4-1) and BD Horizon Fixable Viability Stain 780. Western blot antibodies were purchased from (i) abcam: anti-human CTPS1 (clone EPR8086(B)); (ii) BD Biosciences: anti-human clathrin (clone 23/Clathrin Heavy Chain); (iii) Dako: HRP-labeled goat anti-mouse and swine anti-rabbit immunoglobulins.

### Cell culture and isolation

Cell lines were obtained from the American Type Culture Collection and were maintained at 37°C with 5% CO2. Jurkat, RPMI1788 and Raji-GFP cells were cultured in Roswell Park Memorial Institute 1640 medium (Gibco, Thermo Fisher Scientific) supplemented with 10% fetal bovine serum (FBS, Biowest) and 2 mM L-glutamine (Gibco, Thermo Fisher Scientific). HEK293T cells were cultured in Dulbecco’s Modified Eagle Medium (Gibco, Thermo Fisher Scientific) supplemented with 10% FBS (Biowest) and 1% HEPES (Gibco, Thermo Fisher Scientific). Peripheral blood mononuclear cells (PBMCs) were isolated from buffy coats (Red Cross Belgium) by density gradient centrifugation using Lymphoprep (Alere Technologies AS) and HetaSep (STEMCELL Technologies) to remove red blood cells. Naive CD4^+^ T cells were isolated by negative selection with the EasySep Human Naïve CD4^+^ T Cell Isolation Kit (STEMCELL Technologies) according to manufacturer’s protocol. CD4^+^ and CD8^+^ T cells were isolated by negative selection with the Dynabeads Untouched Human CD4 T Cells Kit and the Dynabeads Untouched Human CD8 T Cells Kit (Invitrogen, Thermo Fisher Scientific) respectively, according to manufacturer’s protocol.

### Plasmids

The pcDNA3.1-hCD4-tGFP-P2A-mCherry construct was cloned by assembly of PCR fragments (New England BioLabs) from the pcDNA3.1 expression vector (Invitrogen, Thermo Fisher Scientific) encoding wild-type hCD4 which was kindly provided by Dr. O. Schwartz (Institut Pasteur, Paris), and the pEGFP-N1 vector (Clontech) containing EGFP-P2A-mCherry, kindly provided by Dr. R. Hegde (MRC, Cambridge). The pcDNA3.1-mCD4 expression vector was generated by cloning full-length mCD4 from a pReceiver-M16 vector, containing mouse CD4-eYFP (GeneCopoeia), into a pcDNA3.1 tGFP-P2A-mCherry vector. The pcDNA3.1-hmCD4-tGFP-P2A-mCherry expression vector was generated by cloning a synthesized gBlock-fragment (IDT) encoding the hCD4-mCD4 sequence into a pcDNA3.1 tGFP-P2A-mCherry vector (Invitrogen, Thermo Fisher Scientific). The other pcDNA3.1-tGFP-P2A-mCherry reporter constructs were cloned by assembly of PCR fragments (New England BioLabs) from different sources: the CD8α reporter construct was generated from a pORF-hCD8α vector purchased from InvivoGen, while the CD25, CD28, CD69, OX40 and 4-1BB reporter constructs were cloned from vectors purchased from Sino Biological. Sequences were confirmed by automated capillary Sanger sequencing (Macrogen Europe).

### Cell transfection

HEK293T cells were plated at 5 × 10^5^ cells/mL in Corning Costar 6-well plates and were transfected with the tGFP-P2A-mCherry constructs 24h after plating. Transfections were done by making use of Lipofectamine 2000 transfection reagent (Invitrogen, Thermo Fisher Scientific). Six hours after transfection, indicated amounts of CADA or 0.1% of DMSO were added. Cells were collected for flow cytometric analysis 24h after transfection.

### Cell viability analysis

Jurkat cells were plated at 1 × 10^5^ cells/mL in Corning Costar 24-well plates in the presence of indicated amounts of CADA or MMF. After 48h, cells were stained with trypan blue and counted with a Vi-CELL cell counter (Beckman Coulter).

### MTS-PES assay

Jurkat cells were plated at 2.5 × 10^5^ cells/mL in Falcon flat-bottom 96-well plates in the presence of indicated amounts of CADA or MMF, or in the presence of corresponding DMSO concentrations. MTS-PES (Promega) was added 48h later and after a 2h incubation period, colorimetric detection was done using the VersaMax microplate reader (Molecular Devices).

### T cell activation by superantigens

Jurkat or naive CD4^+^ T cells were plated at 2.8 × 10^5^ cells/mL in Falcon round-bottom 96-well plates in presence or absence of 10 μM of CADA. After 48h, T cells were activated by adding Staphylococcal enterotoxin E (Toxin Technology) or Staphylococcal enterotoxin B (Sigma-Aldrich)-stimulated Raji-GFP cells at a concentration of 1.2 × 10^6^ cells/mL. Raji cells were labeled with GFP to distinguish them from Jurkat and naive CD4^+^ T cells by flow cytometry. Expression of the early activation marker CD69 was detected by flow cytometry 24h later.

### Mixed lymphocyte reaction

PBMCs (1.2 × 10^6^ cells/mL) were co-incubated with mitomycin C (Sigma-Aldrich)-inactivated RPMI1788 cells (0.45 × 10^6^ cells/mL) in Falcon flat-bottom 96-well plates in the presence of indicated amounts of compounds or antibody and corresponding concentrations of DMSO. At day 5, 0.001 mCi of [^3^H]-thymidine (PerkinElmer) was added per well and 18h later, cells were harvested on Unifilter-96 GF/C plates (PerkinElmer) with the Unifilter-96 Cell Harvester (PerkinElmer). 20 μL of MicroScint-20 (PerkinElmer) was added per filter and counts per minute (cpm) were detected with the MicroBeta device (PerkinElmer). Expression of hCD4 was measured at day 5 by flow cytometry.

### Cell-mediated lympholysis

PBMCs (4.8 × 10^6^ cells/mL) were co-incubated with mitomycin C (Sigma-Aldrich)-inactivated RPMI1788 stimulator cells (1.8 × 10^6^ cells/mL) in Falcon round-bottom 14 mL tubes in presence or absence of CADA for 6 days. After this incubation period, PBMCs were collected and concentrated at 5 × 10^6^ cells/mL. Fresh target RPMI1788 cells were labeled with ^51^Cr (MP Biomedicals), followed by a 4h incubation at 37°C with the PBMCs in a ratio of 50/1 (500,000 effector cells/10,000 target cells per well). To measure spontaneous and maximum release of ^51^Cr, medium or saponin was added to the ^51^Cr-labeled RPMI1788 cells, respectively. After incubation, supernatant was collected and ^51^Cr release was detected using a TopCount gamma counter (Packard Instrument Company). The percentage of specific lysis was calculated by the following formula: % specific lysis = (experimental release – spontaneous release) / (maximum release – spontaneous release) × 100.

### T cell activation by CD3/CD28 beads or phytohemagglutinin

PBMCs were pre-incubated at a concentration of 4 × 10^5^ cells/mL with 10 μM of CADA or 0.1% DMSO during 3 days in Falcon flat-bottom 96-well plates. T cells were activated with Dynabeads Human T-Activator CD3/CD28 (beads/cell ratio of 1/2; Gibco, Thermo Fisher Scientific) or with 4.5 μg/mL phytohemagglutinin (PHA; Sigma-Aldrich) and further incubated with 10 μM of CADA or 0.1% DMSO. At 4h, 1 day, 2 days, 3 days or 4 days after activation, 0.001 mCi of [^3^H]-thymidine (PerkinElmer) was added per well and 22h later, cells were harvested on Unifilter-96 GF/C plates (PerkinElmer) with the Unifilter-96 Cell Harvester (PerkinElmer). 20 μL of MicroScint-20 (PerkinElmer) was added per filter and cpm were detected with the MicroBeta device (PerkinElmer). Expression of CD4, CD8, CD25 and CD28 was measured by flow cytometry just before activation (0h) and 4h, 1 day, 2 days, 3 days or 4 days after activation. Expression of OX40 and 4-1BB was measured by flow cytometry 2 days after activation. Intracellular levels of phosphorylated signal transducer and activator of transcription 5 (pSTAT5) were measured by flow cytometry 2 days after activation with or without an extra stimulation with 25 ng/mL IL-2 (R&D Systems) during 15 min.

### Flow cytometry

Cells were collected and washed in PBS (Gibco, Thermo Fisher Scientific) supplemented with 2% FBS (Biowest). Antibodies were diluted in PBS with 2% FBS and samples were stored in PBS containing 1% formaldehyde (VWR Life Science AMRESCO). For intracellular staining, samples were immediately fixed in PBS with 2% formaldehyde, after which cells were permeabilized using absolute methanol (Biosolve) and stained with antibody. Acquisition of all samples was done on a BD FACSCanto II flow cytometer (BD Biosciences) with BD FACSDiva v8.0.1 software, except for the samples of the tGFP-P2A-mCherry constructs, that were acquired on a BD LSRFortessa flow cytometer (BD Biosciences) with BD FACSDiva v8.0.2 software. Flow cytometric data were analyzed in FlowJo v10.1.

### ELISA and Bio-Plex assay

For detection of soluble CD25 (sCD25), supernatants were collected at 2 days, 3 days or 4 days after activation with CD3/CD28 beads or PHA. The concentration of sCD25 was measured with the Human CD25/IL-2R alpha Quantikine ELISA kit (R&D Systems) according to manufacturer’s protocol. Detection was done using a SpectraMax Microplate Reader (Molecular Devices). For the quantification of the cytokines IL-2, IFN-γ and TNF-α Cytokine Human ProcartaPlex Panel kits (Invitrogen, Thermo Fisher Scientific) were used following manufacturer’s protocol. Supernatant was taken at day 5 post stimulation for the MLR samples and at day 3 for the CD3/CD28 beads- and PHA-activated samples. Detection was done with the Bio-Plex 200 System (Bio-Rad).

### Western blot

PBMCs were lysed in ice-cold Nonidet P-40 buffer (50 mM Tris-HCl (pH 8.0), 150 mM NaCl, 1% Nonidet P-40) supplemented with 100x cOmplete Protease Inhibitor Cocktail (Roche, Sigma-Aldrich) and 250x PMSF Protease Inhibitor (100 mM in dry isopropanol, Thermo Fisher Scientific) and centrifuged at 17,000xg during 10 min. Samples were boiled in reducing 2x Laemmli sample buffer (120 mM Tris-HCl (pH 6.8), 4% sodium dodecyl sulphate, 20% glycerol, 100 mM dithiothreitol, 0.02% bromophenol blue) and were separated on 4-12% Criterion XT Bis-Tris Precast gels (Bio-Rad) using 1x MES buffer (Bio-Rad). After electrophoresis, gels were blotted onto polyvinylidene difluoride membranes with the Trans-Blot Turbo Transfer System (Bio-Rad). Membranes were blocked during 1h with 5% nonfat dried milk in tris-buffered saline with Tween 20 (20 mM Tris-HCl (pH 7.6), 137 mM NaCl, 0.05% Tween 20) and incubated overnight with the primary antibodies at 4°C. The next day, membranes were washed and incubated with the secondary antibodies. Detection was done with a ChemiDoc MP Imaging System (Bio-Rad) using the SuperSignal West Femto Maximum Sensitivity Substrate (Pierce, Thermo Fisher Scientific). Clathrin was used as a control for protein concentration.

### Cell-free in vitro translation and translocation

The Qiagen EasyXpress linear template kit was used to generate full length cDNAs using PCR. PCR products were purified and transcribed *in vitro* using T7 RNA polymerase (RiboMAX system, Promega). All transcripts were translated in rabbit reticulocyte lysate (Promega) in the presence of L-35S-methionine (Perkin Elmer). Translations were performed at 30°C in the presence or absence of ovine pancreatic microsomes and CADA as described elsewhere (Vermeire et al., 2015). Samples were washed with low-salt buffer (80 mM KOAc, 2 mM Mg(OAc)2, 50 mM HEPES pH 7.6) and radiolabeled proteins were isolated by centrifugation for 10 minutes at 21,382×g and 4°C (Hettich 200R centrifuge with 2424-B rotor). The proteins were then separated with SDS-PAGE and detected by phosphor imaging (Cyclone Plus storage phosphor system, Perkin Elmer).

### Statistical analysis

Data were visualized as means ± standard deviation (SD) or as absolute individual data points and were analyzed by making use of the GraphPad Prism v7.0 software. Data were analyzed with multiple t-tests to compare different treatment concentrations to the corresponding control or to compare CADA to DMSO in several stimulation conditions. In case of multiple testing, a Holm-Sidak method was used to correct for multiple comparison. Paired t-tests were used for the comparison of CADA and DMSO for proliferation response, receptor expression and levels of sCD25 at certain time points. P-values bellow 0.05 were considered statistically significant.

## ABBREVIATIONS

CADA: cyclotriazadisulfonamide
CD: cluster of differentiation
CTPS1: cytidine triphosphate synthase 1
ER: endoplasmic reticulum
hCD4: human CD4
IL: interleukin
Lck: lymphocyte C-terminal Src kinase
mCD4: murine CD4
MLR: mixed lymphocyte reaction
MMF: mycophenolate mofetil
PBMC: peripheral blood mononuclear cell
PHA: phytohemagglutinin
pSTAT5: phosphorylated signal transducer and activator of transcription 5
sCD25: soluble CD25
SP: signal peptide

## ACKNOWLEDGEMENTS

We thank Anita Camps, Sandra Claes, Eric Fonteyn, Becky Provinciael, Omer Rutgeerts, Geert Schoofs and Joren Stroobants for their excellent technical assistance. Dr. Thomas W. Bell (UNR, Nevada, USA) is acknowledged for providing CADA compound. We are grateful to Prof. Enno Hartmann and Prof. Kai-Uwe Kalies for providing microsomal membranes. This work was partly supported by the KU Leuven grant no. PF/10/018. BS is a senior clinical investigator of the Research Foundation Flanders (FWO) (1842919N).

## AUTHOR CONTRIBUTIONS

K.V., E.C., B.S. and S.H.-B. conceived experiments; E.C. and E.P. performed experiments; K.V., E.C. and B.S. wrote the manuscript; D.S. secured funding; S.H.-B. and D.S. provided reagents; M.W., B.S. and S.H.-B. provided expertise and feedback.

