## Supplemental figures for "Simultaneous inhibition of human CD4 and 4-1BB biogenesis suppresses cytotoxic T lymphocyte proliferation"

ORCID iD: Kurt Vermeire: 0000-0003-1123-1907

#### SUPPLEMENTARY FIGURES

**Figure 1–figure supplement 1.** CADA has no effect on superantigen-activated lymphocytes. **(A)** Jurkat cells were pre-treated with 10 µM of CADA (green; MFI=13287) for 2 days and subsequently activated with Raji-GFP cells loaded with staphylococcal enterotoxin E (SEE). Cell surface CD69 expression was measured 24h later. Nonactivated cells are represented in blue, activated cells without treatment in red (MFI=20777) and unstained control as a black dotted line. **(B)** Naive $CD4^+$ T cells were pre-treated with 10 µM of CADA (green) for 2 days and subsequently activated with Raji-GFP cells loaded with staphylococcal enterotoxin E (SEE) or staphylococcal enterotoxin B (SEB). CD69 expression was measured 24h later. Nonactivated cells are given in blue, activated cells without treatment in red and unstained control as a black dotted line.

**Figure 1–figure supplement 2.** CADA does not exert cytotoxicity. Jurkat cells were incubated with different concentrations of CADA (red; n=4) or MMF (black; n=4) during 2 days, after which they were stained with trypan blue and counted with a Vi-CELL device. Cell viability is given as percentage of untreated control (mean ± SD).

**Figure 4–figure supplement 1.** CADA suppresses CD4 and CD8 receptor upregulation. PBMCs were treated and collected as described in the legend to Figure 4. CD4 (blue) and CD8 (green) receptor expression analysis was performed on the samples from Figure 4 for the time points 0h (d0), day 1 (d1) and day 4 (d4) post activation. For each donor, the MFI value of the CADA-treated sample (10 µM CADA) is normalized to the corresponding DMSO control (set as 1.00) to calculate the relative receptor expression. Bars are mean ± SD; n=4.

**Figure 6–figure supplement 1.** CADA inhibits surface CD25 upregulation on $CD4^+$ and $CD8^+$ T cells after activation by CD3/CD28 beads or PHA. **(A)** PBMCs were pre-incubated with CADA (10 µM) or DMSO for 3 days, after which they were activated with CD3/CD28 beads. Cellular surface CD25 expression was measured on gated $CD4^+$ (left panel) and $CD8^+$ (right panel) T cells by flow cytometry just before activation (0h) and 4h, 1 day, 2 days, 3 days or 4 days post activation. Mean fluorescence intensity (MFI) of CD25 expression is shown for 4 donors of PBMCs with DMSO-treated samples as a dotted line and CADA-treated cells as a purple solid line. The same donors from Figure 6 were used. **(B)** CD25 receptor expression analysis was performed on the samples from (A) and from Figure 6 for the time points 0h (d0), day 1 (d1) and day 4 (d4) post activation. For each donor, the MFI value of the CADA-treated sample (10 μM CADA) is normalized to the corresponding DMSO control (set as 1.00) to calculate the relative receptor expression. Bars are mean ± SD; n=4.

**Figure 6–figure supplement 2.** CADA reduces the levels of soluble CD25 protein. PBMCs were pre-incubated with 10 μM of CADA or DMSO during 3 days, after which they were activated by CD3/CD28 beads (left) or PHA (right). Supernatant was taken 2, 3 or 4 days post activation and sCD25 levels were measured by ELISA. Data are shown for 4 donors of PBMCs with DMSO-treated samples as open symbols and CADA-treated samples as solid purple symbols. A paired t-test was performed to compare CADA to DMSO with *$p<0.05$.

**Figure 7–figure supplement 1.** CADA reduces the upregulation of CD28 on lymphocytes after activation by CD3/CD28 beads or PHA. **(A and B)** PBMCs were pre-incubated with 10 μM of CADA or DMSO during 3 days, after which they were activated by CD3/CD28 beads or PHA. **(A)** Cell surface CD28 expression was measured on gated $CD4^+$ (top panels) and $CD8^+$ (bottom panels) T cells by flow cytometry just before activation (0h) and 4h, 1 day, 2 days, 3 days or 4 days post activation. Mean fluorescence intensity (MFI) of CD28 expression is shown for 4 donors of PBMCs (indicated in different color) with DMSO-treated samples as a full line and CADA-treated cells as a dotted line. **(B)** For each donor, the MFI value of the CADA-treated sample is normalized to the corresponding DMSO control (set as 1.00) to calculate the relative CD28 receptor expression. Bars are mean ± SD; n = 4. Multiple t-tests were performed to compare CADA to DMSO for each condition with *$p<0.05$ and with Holm-Sidak method as correction for multiple comparison.

**Figure 9–figure supplement 1.** CADA reversibly down-modulates hCD4. HEK293T Cells were transfected with hCD4tGFP-2A-RFP and given DMSO (CTR) or treated with CADA for 72h. In parallel, CADA-treatment was terminated after 24h (CADA wash). These cells were washed profoundly and given control medium for the duration of the experiment. At the indicated time points, cells were collected and tGFP (representing human CD4 levels) was measured by flow cytometry. The average MFI of tGFP is shown (mean ± SD; n=2). Of note is that the SD of the control samples (grey curve) and CADA samples (green curve) is too small to be visible on the graph.

**Figure 9–figure supplement 2.** CADA differentially affects the protein expression levels of co-stimulatory receptors in transfected cells. HEK293T cells were transiently transfected with different receptor constructs (as shown in Figure 9A) and incubated with CADA. Cellular expression of each receptor-tGFP-2A-RFP was determined by measuring tGFP levels by flow cytometry. Receptor levels in CADA-treated samples are normalized to the corresponding DMSO control. Values are mean ± SD; n ≥ 3. For each construct, a four parameter dose-response curve for CADA was fitted to data from at least three replicate experiments.

**Figure 10–figure supplement 1.** Cell free *in vitro* translation/translocation assay to study the co-translational translocation of proteins. **(A)** Schematic representation of translating ribosomes targeting to the ER membrane. When the signal peptide (SP) is emerging from the translating ribosome (1), the ribosome-nascent chain complex is targeted to the ER membrane and will dock onto the Sec61 translocon. The SP is then inserted in the translocon and makes a looped structure with its N-terminus facing the cytosol (2). The signal peptidase, located at the lumenal side of the ER membrane, will cleave the SP from the pre-protein (3). Next, the oligosaccharyltransferase (OST), also located at the lumenal side of the ER membrane, will add glycans to the protein (4). Finally, a lateral gate in the translocon facilitates the integration of hydrophobic transmembrane protein segments into the lipid bilayer for the accommodation of integral membrane proteins (5). **(B)** Schematic overview of the protocol used in the translation/translocation assay.

### Figure 1 - figure supplement 1

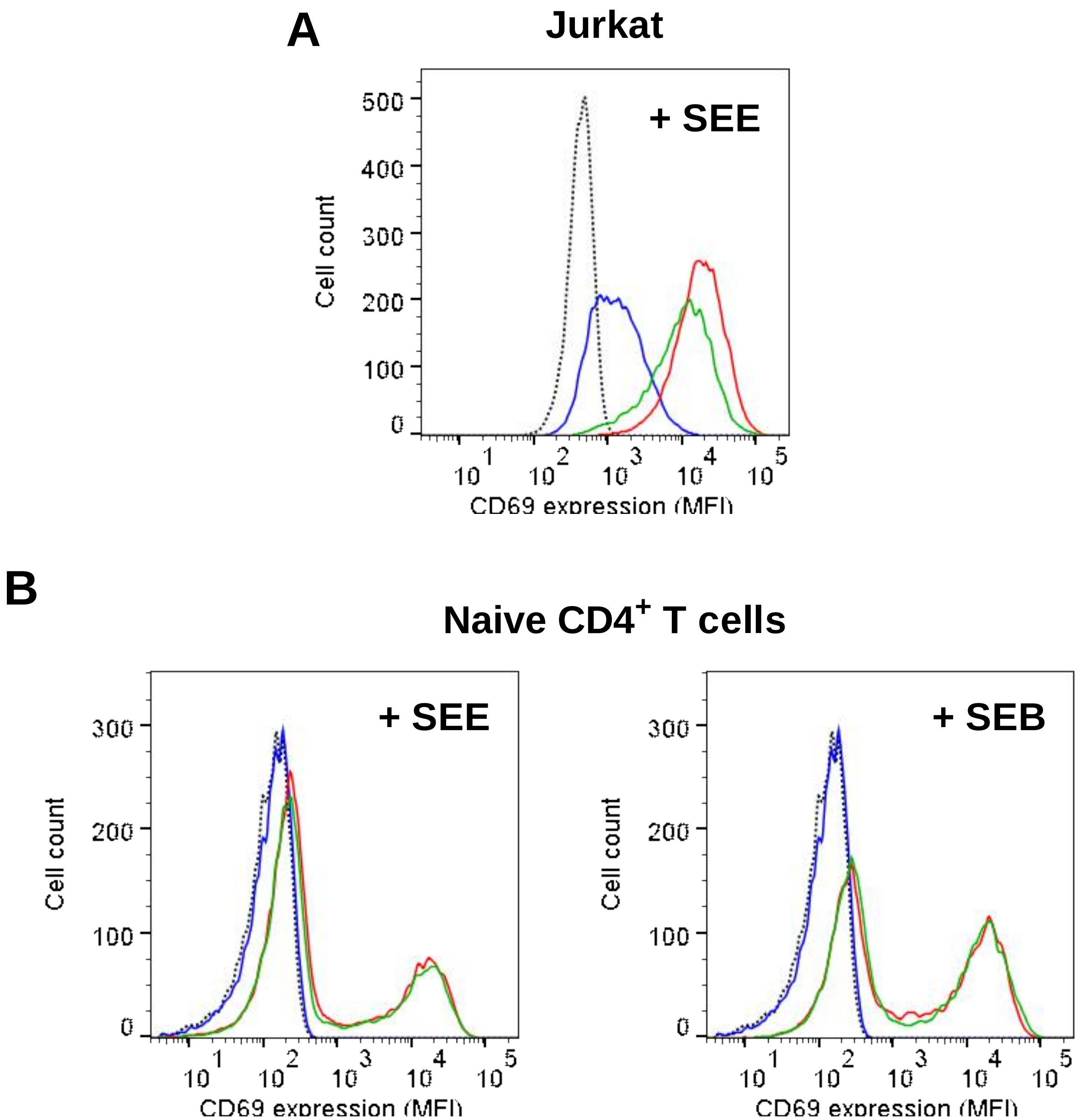


Claeys et al.

### Figure 1 - figure supplement 2

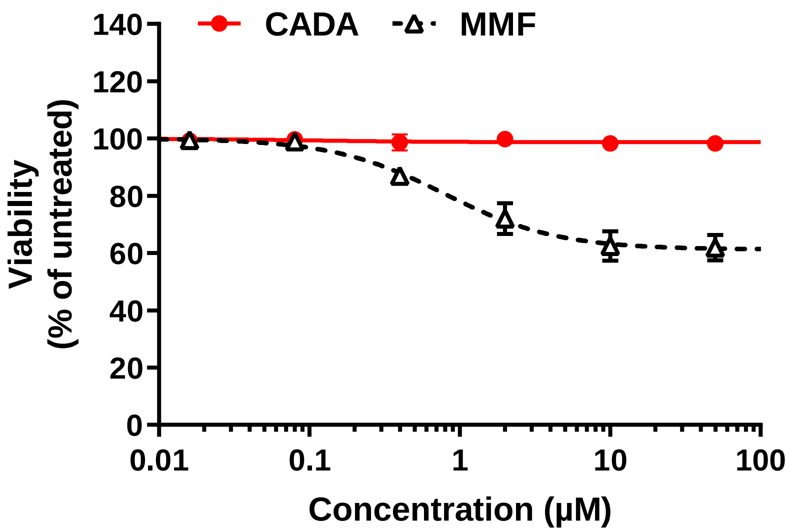


Claeys et al.

### Figure 4 - figure supplement 1

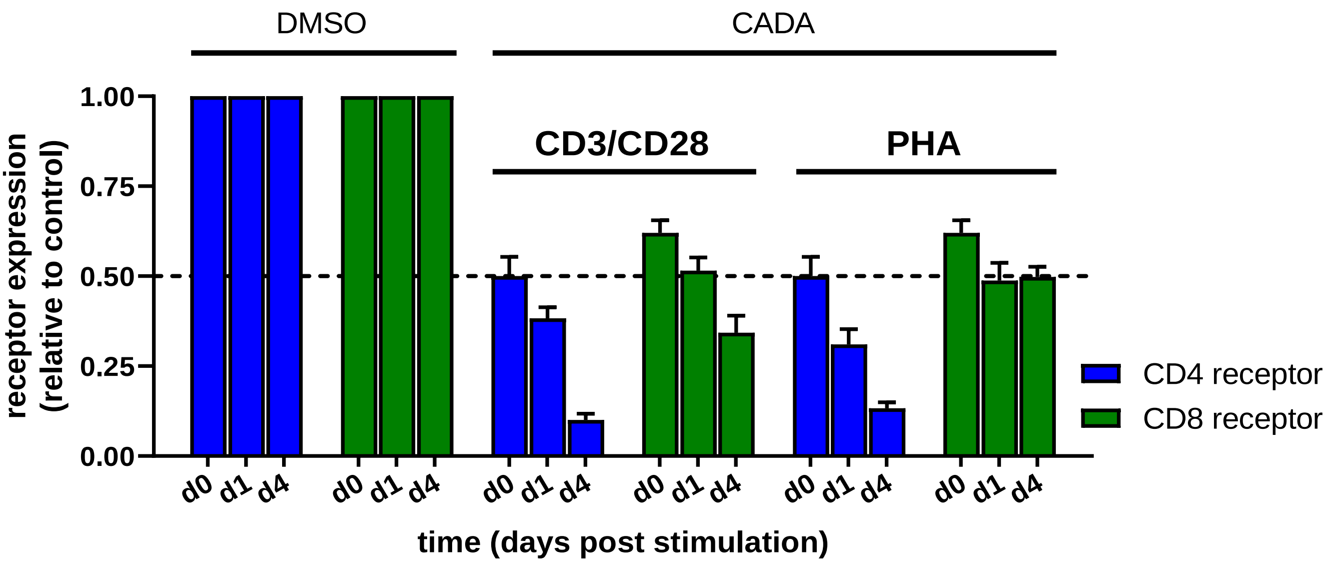


Claeys et al.

### Figure 6 - figure supplement 1

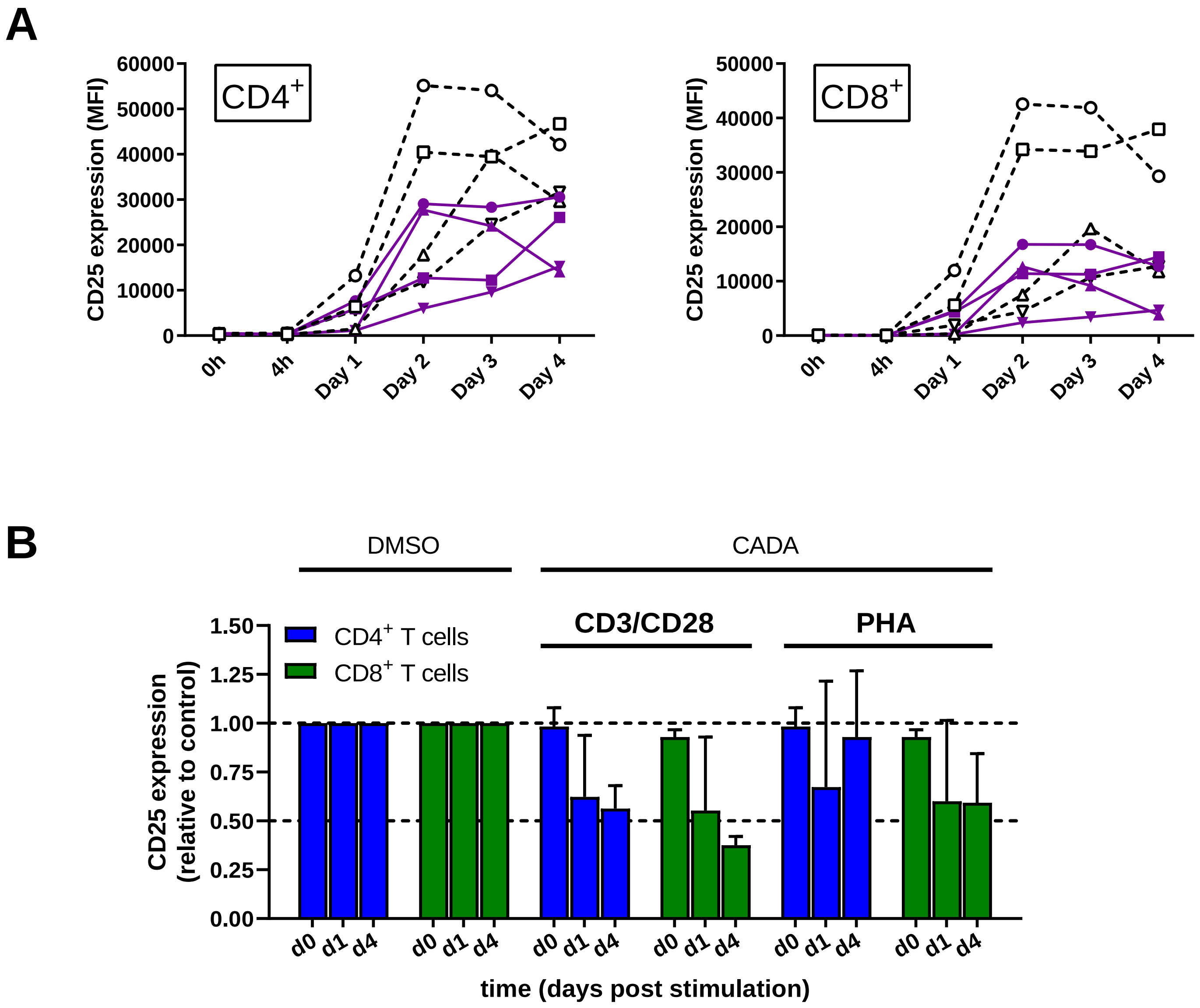


Claeys et al.

### Figure 6 - figure supplement 2

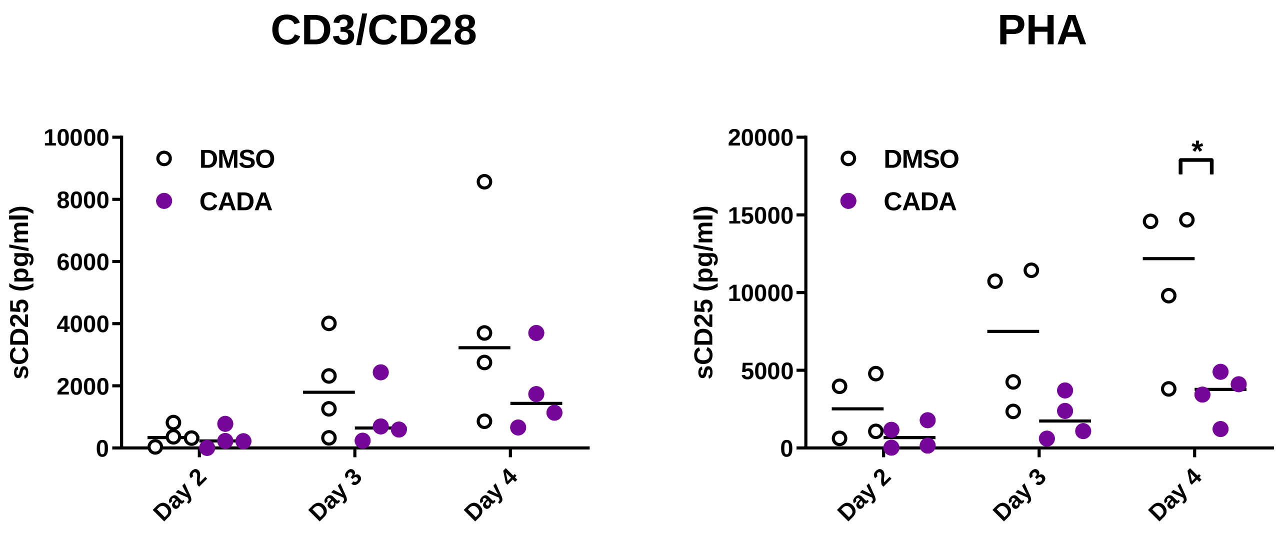


Claeys et al.

### Figure 7 - figure supplement 1

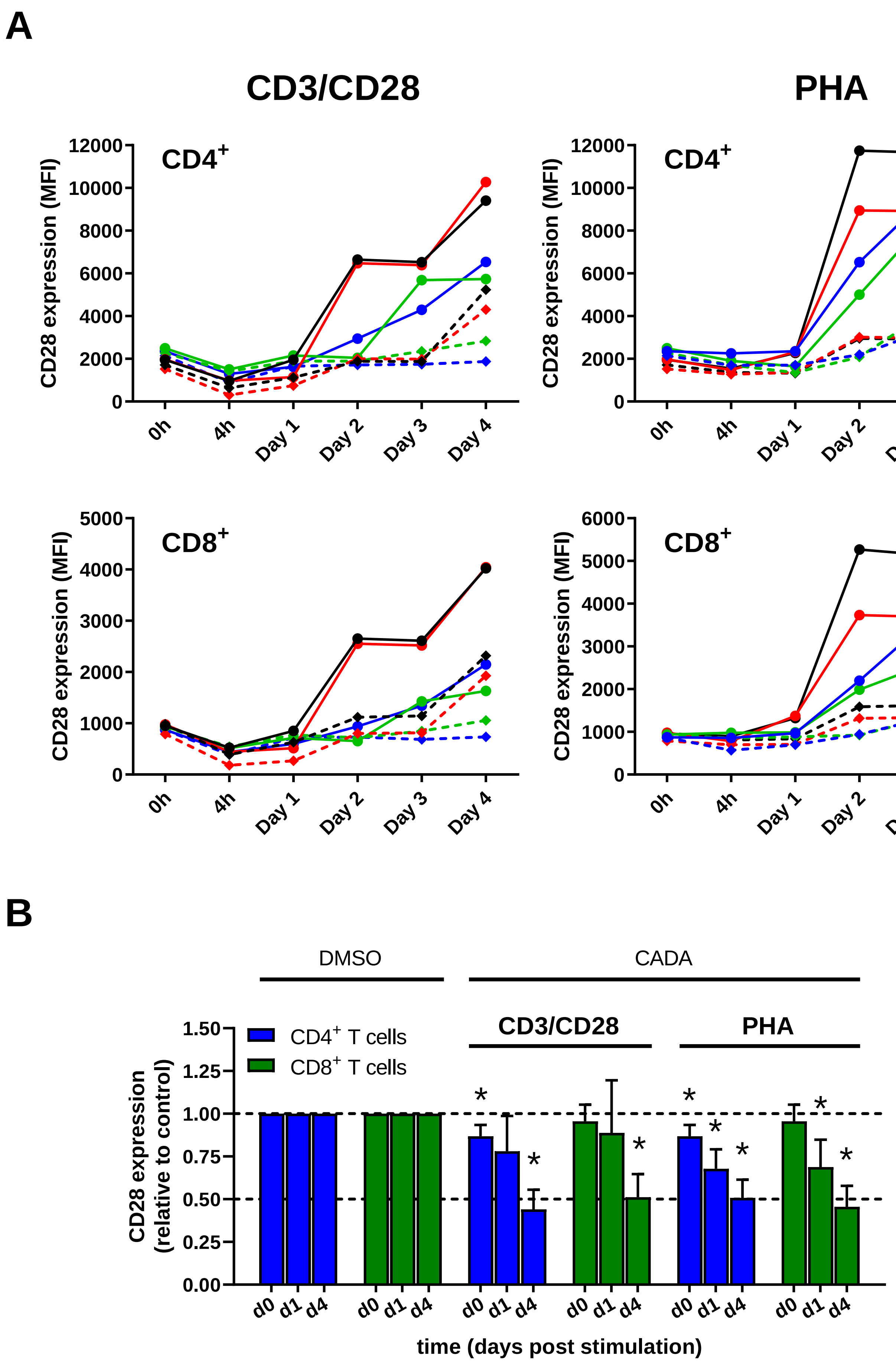

### Figure 9 - figure supplement 1

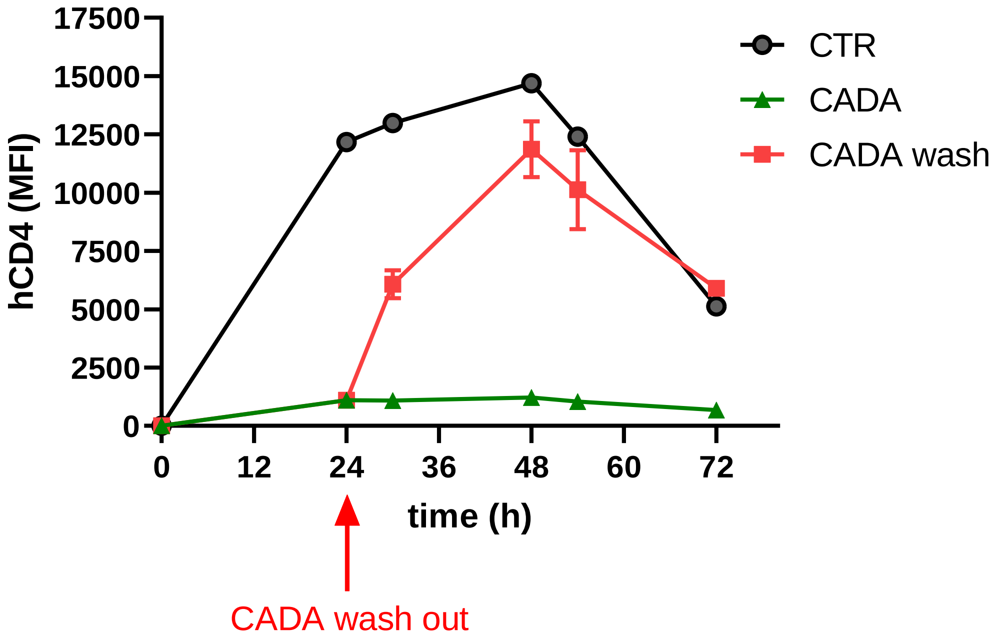

### Figure 9 - figure supplement 2

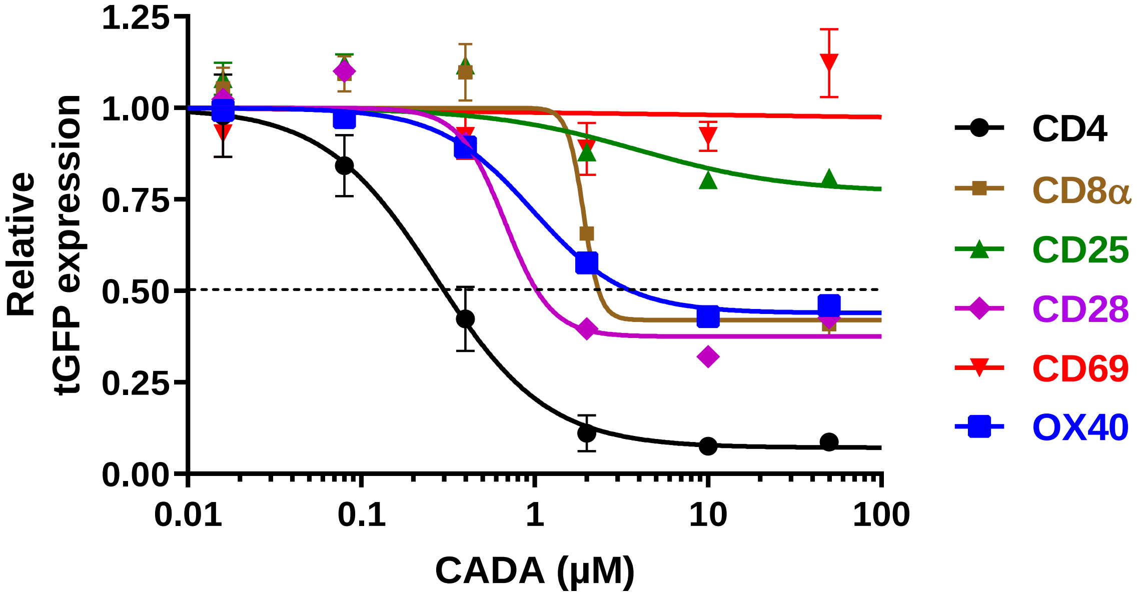

### Figure 10 - figure supplement 1

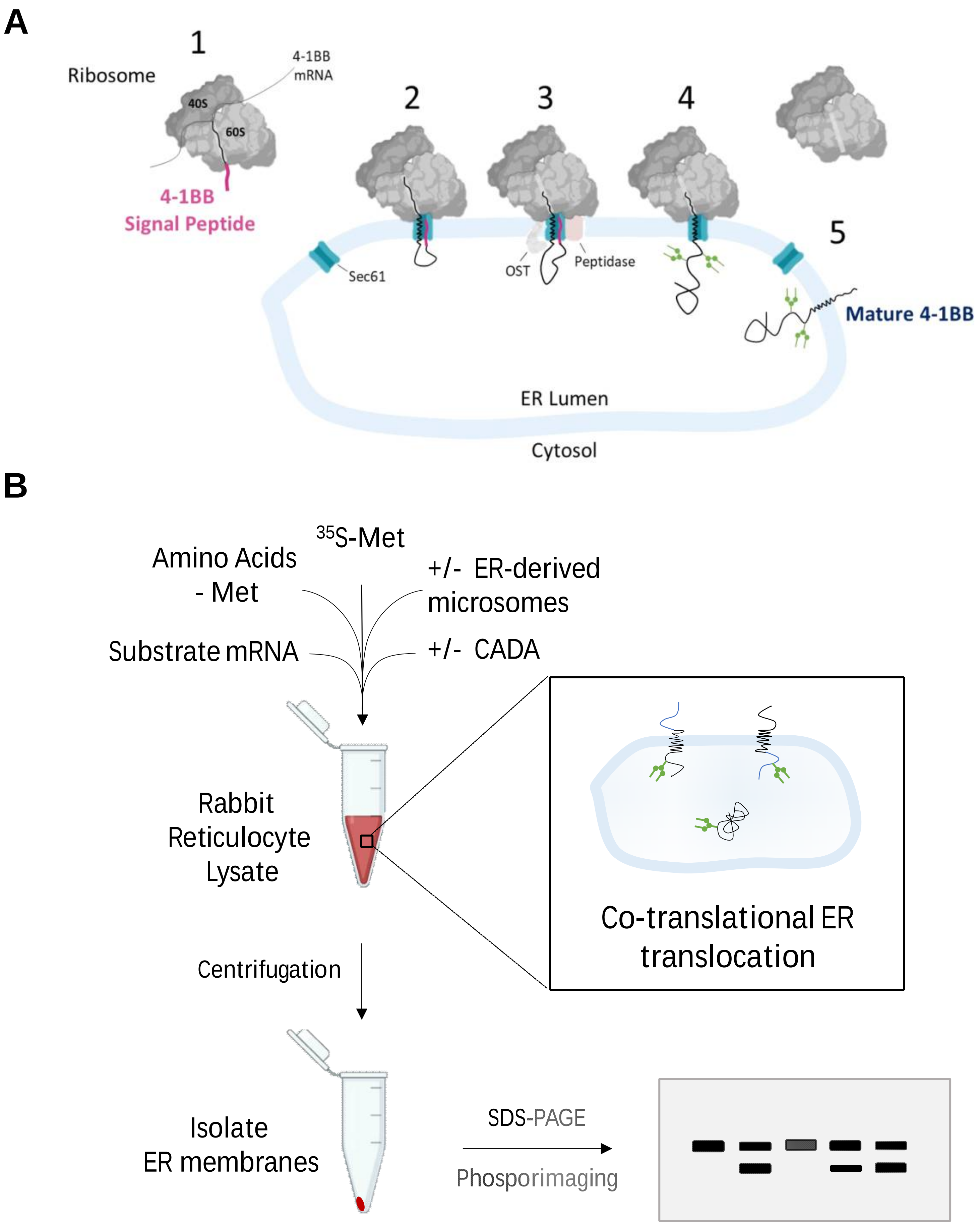
